# The nutrient-sensing Rag-GTPase complex in B cells controls humoral immunity via TFEB/TFE3-dependent mitochondrial fitness

**DOI:** 10.1101/2024.02.26.582122

**Authors:** Xingxing Zhu, Yue Wu, Yanfeng Li, Xian Zhou, Jens O. Watzlawik, Yin Maggie Chen, Ariel L. Raybuck, Daniel Billadeau, Virginia Shapiro, Wolfdieter Springer, Jie Sun, Mark R. Boothby, Hu Zeng

## Abstract

During the humoral immune response, B cells undergo rapid metabolic reprogramming with a high demand for nutrients, which are vital to sustain the formation of the germinal centers (GCs). Rag-GTPases sense amino acid availability to modulate the mechanistic target of rapamycin complex 1 (mTORC1) pathway and suppress transcription factor EB (TFEB) and transcription factor enhancer 3 (TFE3), members of the microphthalmia (MiT/TFE) family of HLH-leucine zipper transcription factors. However, how Rag-GTPases coordinate amino acid sensing, mTORC1 activation, and TFEB/TFE3 activity in humoral immunity remains undefined. Here, we show that B cell-intrinsic Rag-GTPases are critical for the development and activation of B cells. RagA/RagB deficient B cells fail to form GCs, produce antibodies, and generate plasmablasts in both T-dependent (TD) and T-independent (TI) humoral immune responses. Deletion of RagA/RagB in GC B cells leads to abnormal dark zone (DZ) to light zone (LZ) ratio and reduced affinity maturation. Mechanistically, the Rag-GTPase complex constrains TFEB/TFE3 activity to prevent mitophagy dysregulation and maintain mitochondrial fitness in B cells, which are independent of canonical mTORC1 activation. TFEB/TFE3 deletion restores B cell development, GC formation in Peyer’s patches and TI humoral immunity, but not TD humoral immunity in the absence of Rag-GTPases. Collectively, our data establish Rag-GTPase-TFEB/TFE3 pathway as an mTORC1 independent mechanism to coordinating nutrient sensing and mitochondrial metabolism in B cells.

**One sentence summary:** Rag-GTPases restrain TFEB/TFE3 to prevent abnormal mitophagy, maintain mitochondrial fitness, and support B cell development, Peyer’s patch germinal center formation, and T-independent humoral immunity.

## Introduction

During infection or immunization, B lymphocytes can be activated in a T cell-dependent (TD) or independent (TI) manner, differentiate into plasmablast cells or form germinal center (GC) and subsequently differentiate into memory B cells and plasma cells (*1–4*), which produce different isotypes of antibodies (*5*). There is increasing evidence that metabolic programming underpins B cell development, quiescence, activation, and differentiation (*6–8*). Specifically, glucose metabolism is vital to support the B cell function like early B cell development in bone marrow (BM), affinity maturation of GC B cells, and lymphomagenesis (*9–11*). Lactate dehydrogenase dependent glycolysis is dispensable for TI responses, but critical for TD responses, highlighting divergent metabolic requirements between TD and TI responses (*10*). Mitochondria oxidative phosphorylation, amplified by CD40L or TLR ligand engagement, supports B cell survival and differentiation (*12–15*). Nutrients modulate cellular metabolic reprogramming, and subsequently affect immune responses (*16–18*). Consequently, malnutrition, including Kwashiorkor, a form of severe protein inadequacy, is associated with small Peyer’s patches (PPs) and GC, fewer antibody-producing cells and increased susceptibility to infection (*19, 20*). A potential mechanism through which amino acid controls immunity is the tuning of mTORC1 activation (*16*). Yet, how B cells sense nutrients to coordinate their mitochondrial metabolism, and how metabolic pathways support B cell development and responses to TD and TI antigens, remains poorly defined.

Rag-GTPases are the key amino acid sensors that relay amino acid availability to modulate mTORC1 and suppress TFEB transcription factor (*21–23*). As small GTPases, Rags are obligate heterodimers, configured such that RagA or RagB is bound to RagC or RagD. RagA is required for amino acid-dependent mTORC1 activation *in vitro* (*21, 23*). Gain of function mutations in Rag-GTPases lead to overactivation of mTORC1 in B cells, demonstrating that Rag-GTPases are sufficient for mTORC1 activation (*24, 25*). Yet, there is controversy regarding the necessity of Rag-GTPases for activation of canonical mTORC1 (*26–30*). In fact, Rag-GTPases can even suppress mTORC1 signaling because RagA/RagB deficient macrophages exhibit highly elevated mTORC1 activity, indicating a cell type specific, context dependent relationship between Rag-GTPases and mTORC1 (*31*). Recent studies have demonstrated that Rag-GTPases cooperate with Rheb to activate mTORC1 to support Treg functions (*32–34*). However, the mechanisms through which Rag-GTPases coordinate mTORC1 and TFEB to regulate humoral immunity are currently unknown.

MiTF/TFE family members, including MITF, TFEB, TFE3, and TFEC, are basic helix-loop-helix leucine zipper (bHLH-Zip) transcription factors. They share a similar structure and often express together (*35*). Among them, TFEB and TFE3 regulate a similar set of genes involved in lysosomal biogenesis, lipid metabolism, autophagy, and stress response (*36–38*). They can undergo cytoplasm-to-nucleus shuttling in response to different nutrition statuses, which is governed by the phosphorylation through kinases such as mTORC1, ERK, and GSK3 (*39–42*). TFEB/TFE3 can promote autophagy and production of proinflammatory cytokine in macrophage *in vitro*, but it can also enhance mTORC1 activation in tumor-associated macrophages (*31, 43, 44*). In adaptive immunity, TFEB/TFE3 support CD40 ligand expression on T cells, maintains regulatory T cell functions and may prevent B cell senescence during aging (*45–47*). Their B cell intrinsic functions in cellular metabolism and humoral immune response have not been explored.

Here, we took a genetic approach to dissect the contributions of Rag-GTPases and mTORC1 to B cell development and function *in vivo*. Our study shows that acute deletion of either RagA/RagB or Raptor blocks early B cell development. Furthermore, B cell intrinsic Rag-GTPases are critically required for GC formation in PPs, dark zone (DZ) and light zone (LZ) distribution in GC, TD and TI antigen immunization-induced antibody responses, but largely dispensable for mTORC1 activity. Mechanistically, inhibition of TFEB/TFE3 activity is required for optimal B cell activation. Rag-GTPases suppress TFEB/TFE3 activity, largely independent of canonical mTORC1 activity, thereby downregulating lysosomal function and preventing abnormal mitophagy to maintain the mitochondrial fitness in B cells. Deletion of TFEB/TFE3 restores mitochondrial functions, humoral immunity towards TI, but not TD, antigen immunization, and GC formation in PPs in the absence of Rag-GTPases. Collectively, our results demonstrate that the amino acid sensing complex Rag-GTPases maintains mitochondrial fitness by suppressing TFEB/TFE3 activity, independent of mTORC1, in B cells to support the humoral immune responses.

## Results

### Amino acids modulates mTORC1 independent of Rag-GTPases

Amino acids (AAs) are the key nutrients engaging Rag-GTPases and mTORC1 signaling(*40*). To evaluate the impact of AAs on B cell activation and mTORC1 activity, we stimulated B cells in full AAs, essential AAs (EAA) or no AA condition. Among the three conditions, full AA induced the largest cell size, highest expression of CD86, amino acid transporter CD98, and phosphorylation of ribosomal protein S6 (p-S6) and eukaryotic initiation factor 4E-binding protein 1 (p-4EBP1), markers for mTORC1 activation, while B cells in no AA condition had the lowest expression of these markers (Fig. 1A). When we activated B cells in the presence of a titrated concentration of AAs, we observed that AAs promote B activation in a concentration dependent manner (Fig. 1B). Taken together, these data suggest that AAs can fine-tune B cell activation and mTORC1 activity.

**Figure 1.**
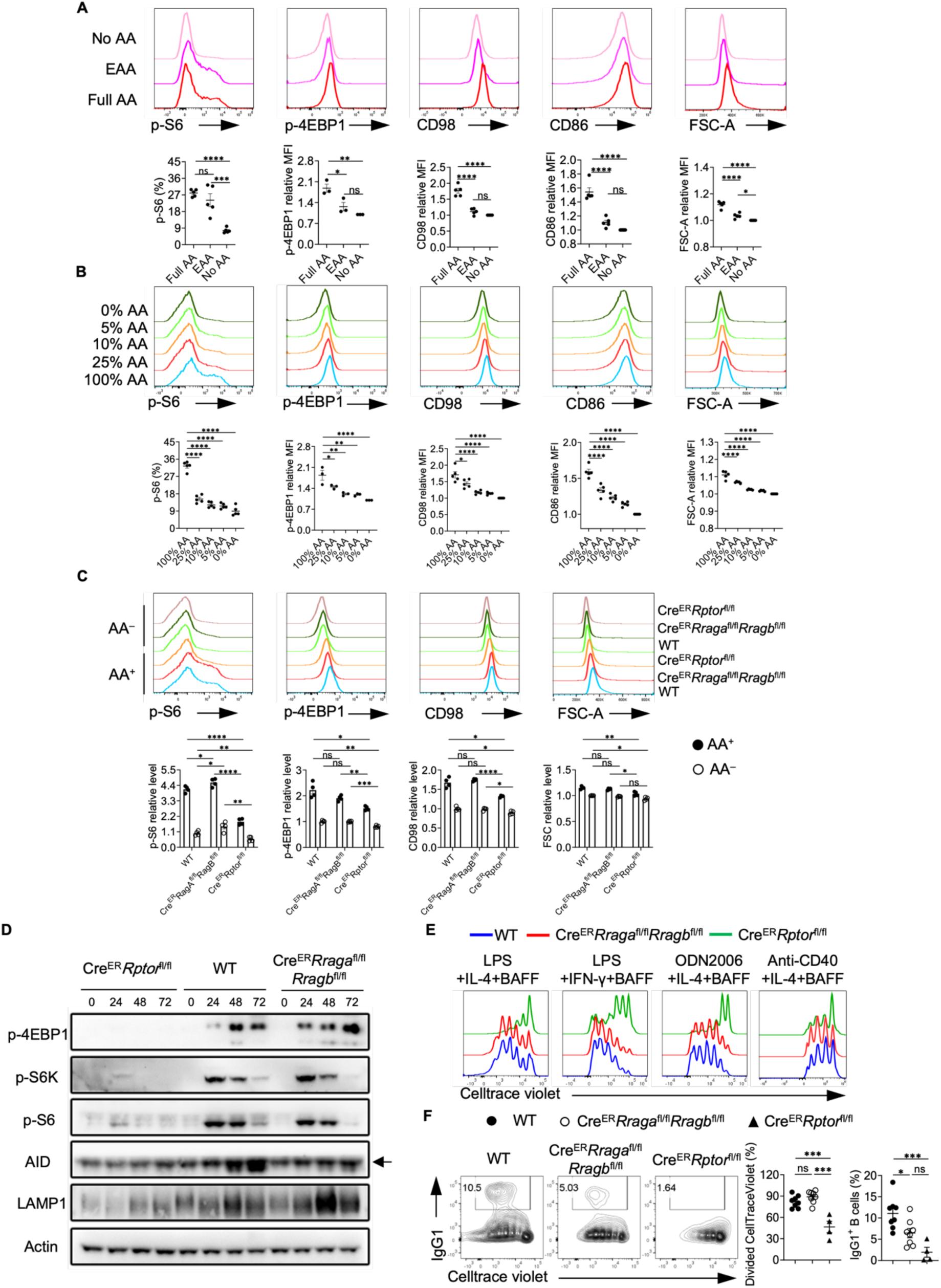
Amino acids modulate mTORC1 independent of Rag-GTPases. (A) B cells were purified and cultured in no amino acids (No AA; n = 5) medium, essential amino acids (EAA; n = 5) medium, or full amino acids (Full AA; n = 5) medium with LPS/IL-4/BAFF overnight. CD98, CD86, p-4EBP1, p-S6 and FSC-A levels were measured by flow cytometry. (B) B cells were cultured in the medium with the indicated concentrations of amino acids with LPS/IL-4/BAFF overnight. CD98, CD86, p-4EBP1, p-S6 and FSC-A levels were measured by flow cytometry. N = 3-5. (C-F) Tamoxifen was administered to animals intraperitoneally daily for 4 consecutive days. Splenic B cells were purified 7 days after the last tamoxifen injection. (C) B cells were stimulated with LPS/IL-4/BAFF overnight in full AA or no AA medium. p-4EBP1, p-S6, CD98 and FSC-A levels were measured by flow cytometry. Cre^ER^ control (WT) mice (n = 4), Cre^ER^ *Rraga*^fl/fl^ *Rragb*^fl/fl^ mice (n = 4), Cre^ER^ *Rptor*^fl/fl^ mice (n = 4). (D) B cells were stimulated with LPS/IL-4/BAFF in full AA medium for the indicated time, and cell lysates were prepared for immunoblotting to detect p-4EBP1, p-S6, p-S6K, AID and LAMP1. β-actin was used as the loading control. Arrow indicates non-specific bands. (E) B cells were labeled with CellTrace violet (CTV) and stimulated with LPS/IL-4/BAFF, LPS/IFN-γ/BAFF, ODN2006/IL-4/BAFF, or Anti-CD40/IL-4/BAFF in complete RPMI1640 medium for 72 h. CTV dilution was measured by flow cytometry. (F) B cells were stimulated with LPS/IL-4/BAFF for 72 h, and IgG1 expression was examined by flow cytometry. Right, summary of the percentages of divided cells and IgG1^+^ B cells. WT mice (n = 8), *Cre^ER^ Rraga*^fl/fl^ *Rragb*^fl/fl^ mice (n = 8), *Cre^ER^ Rptor*^fl/fl^ mice (n = 4). Error bars represent mean ± SEM. ns, not significant. *p < 0.05, **p < 0.01, ***p < 0.001, and ****p < 0.0001, one-way ANOVA (A, B and F), or two-way ANOVA (C).

Given the contention regarding the link between Rag-GTPases and mTORC1 (*26–30*), we directly compared the phenotypes between Rag-GTPase deficiency and mTORC1 deficiency in B cells. RagA and RagB or Raptor were acutely deleted by tamoxifen injection into Cre^ER^*Rraga*^fl/fl^*Rragb*^fl/fl^ mice or Cre^ER^*Rptor*^fl/fl^ mice, which did not significantly alter mature B cell compartment in spleens, including follicular B cells and marginal zone B cells (Fig. S1A-S1B). Raptor deficient splenic B cells had diminished p-S6 and p-4EBP1 levels, reduced CD98 expression and cell size in either full AA medium or no AA medium (Fig. 1C). In contrast, RagA/RagB deficient B cells did not exhibit reduction of any of these parameters in full AA medium. However, under no AA condition, all these parameters reduced to a largely comparable level between 3 genotypes (Fig. 1C), suggesting a Rag-GTPase independent but AA dependent regulation of mTORC1 and initial B cell activation. Immunoblot analysis on B cells activated in full AA medium at different time points confirmed that Raptor deficiency nearly abolished p-4EBP1, p-S6K and p-S6, which were largely intact in the absence of RagA/RagB (Fig. 1D). Consistent with previous publications (*16, 48*), Raptor deficient B cells exhibited substantially reduced proliferation upon various antigenic stimulations. In contrast, RagA/RagB deletion did not affect the proliferation of B cells (Fig. 1E). However, the class switch to IgG1-producing B cells induced by LPS was reduced in both Rag-GTPases or Raptor deficient B cells (Fig. 1F), suggesting a proliferation independent class switch defect in RagA/RagB deficient B cells. Consistent with the reduced IgG1 expression, both Raptor and RagA/RagB deficient B cells showed reduced AID induction (Fig. 1D). Thus, amino acids modulate mTORC1 independent of Rag-GTPases. Rag-GTPases can regulate B cell class switching independent of canonical mTORC1 activity *in vitro*.

### Acute deletion of Rag-GTPases blocks early B cell development and abolishes germinal center B cells in Peyer’s patches

Constitutive deletion of Raptor blocks early B cell development at pro-B to pre-B transition(*8, 49*). While acute deletion of Raptor or RagA/RagB did not affect mature B cells in spleens, we examined the potential impacts on early B cell development. Tamoxifen injected Cre^ER^*Rraga*^fl/fl^*Rragb*^fl/fl^ mice had relatively normal frequencies but reduced absolute numbers of B220^+^CD43^+^IgM^−^ B cell precursors, which were largely intact in tamoxifen injected Cre^ER^*Rptor*^fl/fl^ mice (Fig. 2A-2B). Within B cell precursors, the frequencies of fraction B and fraction C/C’ were significantly reduced in the absence of either Rag-GTPases or mTORC1 (Fig. 2C-2D). Later stage B cells, including B220^+^CD43^−^ pre-B cells/immature B cells and B220^hi^CD43^−^ circulating mature B cells, were mostly reduced in the absence of RagA/RagB (Fig. 2A). In contrast, Raptor deficiency significantly reduced pre-B cells/immature B cells, but increased circulating mature B cells (Fig. 2B). These data indicate that acute loss of either Rag-GTPases or mTORC1 reduces frequencies of fraction B and fraction C/C’ precursors. We did not observe any developmental defects in Cre^ER^*Rragb*^fl/fl^ mice (Fig. S1C-S1D). Thus, both Rag-GTPases and mTORC1 are critically required for early B cell development, but they may have different functions in the maintenance of B220^+^CD43^+^IgM^−^ B cell precursors and circulating mature B cells. In addition, RagA and RagB have redundant functions in B cell development.

**Figure 2.**
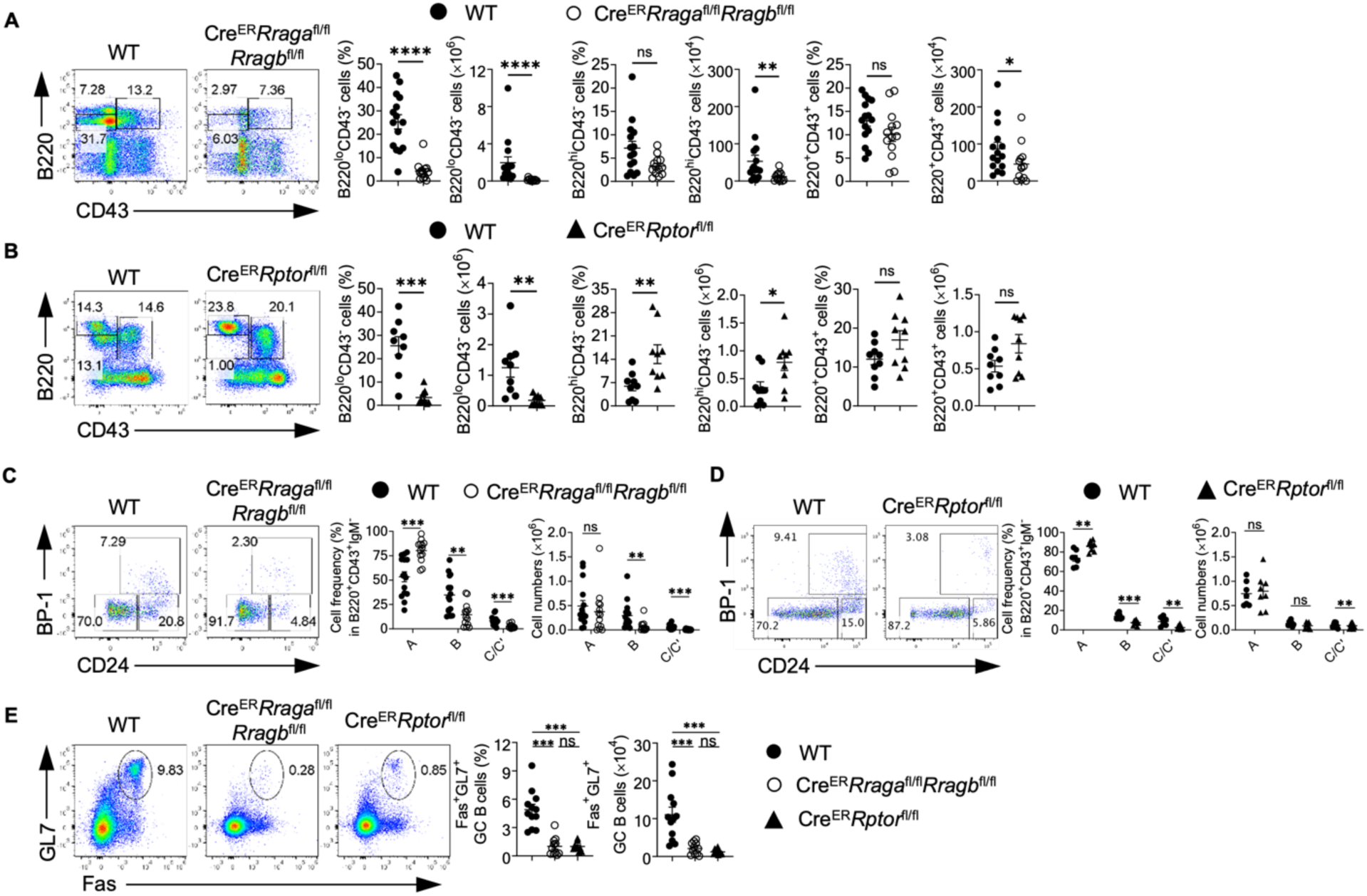
Acute deletion of RagA/RagB blocks early B cell development and abolishes germinal center B cells in Peyer’s patches. (A-E) Tamoxifen was administered to animals intraperitoneally daily for 4 consecutive days. Mice were sacrificed and analyzed 7 days after the last injection. (A) Representative flow plots of B220 and CD43 expression in bone marrow (BM) lymphocytes from WT (n = 15) and *Cre^ER^ Rraga*^fl/fl^ *Rragb*^fl/fl^ (n = 13) mice. Right, summaries of the percentages and numbers of B220^lo^ CD43^−^, B220^hl^ CD43^−^ and B220^+^ CD43^+^ cells. (B) Representative flow plots of B220 and CD43 expression in BM lymphocytes from WT (n = 9) and *Cre^ER^ Rptor*^fl/fl^ (n = 9) mice. Right, summaries of the percentages and numbers of B220^lo^ CD43^−^, B220^hl^ CD43^−^ and B220^+^ CD43^−^ cells. (C) Representative flow plots of BP-1 and CD24 expression in BM B220^+^ CD43^+^ IgM^−^ B cell precursors from WT (n = 15) and *Cre^ER^ Rraga*^fl/fl^ *Rragb*^fl/fl^ (n = 13) mice. Right, summary of the percentages and numbers of fraction A (CD24^−^ BP-1^−^), fraction B (CD24^+^ BP-1^−^), and fraction C/C′ (CD24^+^ BP-1^+^) cells. (D) Representative flow plots of BP-1 and CD24 expression in BM B220^+^ CD43^+^ IgM^−^ B cell precursors from WT (n = 7) and *Cre^ER^ Rptor*^fl/fl^ (n = 8) mice. Right, summary of the percentages and numbers of fraction A (CD24^−^ BP-1^−^), fraction B (CD24^+^ BP-1^−^), and fraction C/C′ (CD24^+^ BP-1^+^) cells. (E) Representative flow plots of GL-7 and Fas expression in lymphocytes from the Peyer’s patches of WT (n = 12), *Cre^ER^ Rraga*^fl/fl^ *Rragb*^fl/fl^ (n = 11), and *Cre^ER^ Rptor*^fl/fl^ (n = 9) mice. Right, summary of the percentages and numbers of GC (GL-7 Fas) B cells. Data in graphs represent mean ± SEM. ns, not significant. *p < 0.05, **p < 0.01, ***p < 0.001, and ****p < 0.0001, two-tailed Student’s t test (A and B), one-way ANOVA (E), two-way ANOVA (C and D).

Acute deletion of either Rag-GTPases or mTORC1 preserves naïve mature CD4^+^ T and B cells in spleens (Fig. S1A-S1B). However, we found that the frequencies and absolute numbers of GC B cells significantly decreased in PPs of both Cre^ER^*Rraga*^fl/fl^*Rragb*^fl/fl^ mice and Cre^ER^*Rptor*^fl/fl^ mice following tamoxifen injection (Fig. 2E), indicating that Rag-GTPases and mTORC1 are vital to the formation of GC in PPs. In all cases, RagB deficiency did not affect mature B cell numbers (Fig. S1E) or GC formation in PPs (Fig. S1F), again demonstrating a redundancy between RagA and RagB.

To study whether the above defects are B cell-intrinsic, we reconstituted lethally irradiated CD45.1^+^ congenic mice with a 4:1 mixture of bone-marrow (BM) recovered from B cell deficient (μMT) mice and either WT (Cre^ER^) or Cre^ER^*Rraga*^fl/fl^*Rragb*^fl/fl^ mice (*50, 51*). Thus, tamoxifen administration achieved acute B cell specific deletion of RagA/RagB. Tamoxifen injected μMT:Cre^ER^*Rraga*^fl/fl^*Rragb*^fl/fl^ mice exhibited reduced B220^lo^CD43^−^ pre-B/immature B cells (Fig. S2A), reduced frequencies of fraction B and C/C’ B cells (Fig. S2B), intact follicular and MZ B cells (Fig. S2C), and greatly reduced GC B cells in PPs (Fig. S2D). Taken together, B cell-intrinsic Rag-GTPases are critically required for early B cell development and GC formation in PPs.

### Rag-GTPases are critically required for GC formation and antibody production independent of mTORC1

To explore the function of B cell-specific Rag-GTPases following T-dependent immune challenge, we immunized tamoxifen injected μMT:Cre^ER^ (WT), μMT:Cre^ER^*Rraga*^fl/fl^*Rragb*^fl/fl^ or μMT:Cre^ER^*Rptor*^fl/fl^ chimera mice with NP-OVA precipitated in alum. Loss of either Rag-GTPases or mTORC1 led to profound loss of GC formation (Fig. 3A, Fig. S3A) and reduced generation of antigen specific B cells in both spleen and mesenteric lymph node (mLN) (Fig. S3B-S3C). Such phenotypes were confirmed with immunofluorescence (Fig. S3D). Consistent with the reduction of GC B cells, expression of Bcl6, a transcriptional factor critical for GC reaction (*52*), was reduced in both mouse strains (Fig. S3E). GC is compartmented into LZ and DZ. DZ B cells are highly proliferative and undergo somatic hypermutation, while LZ B cells undergo selection and affinity maturation (*53*). The ratio of DZ/LZ increased in the remaining GC B cells from μMT:Cre^ER^*Rraga*^fl/fl^*Rragb*^fl/fl^ mice compared to that in control mice, however, the ratio stayed unchanged in the absence of Raptor (Fig. 3B). Furthermore, plasmablast response in spleen was significantly compromised in both knockout strains, with RagA/RagB deficiency effecting a stronger reduction than mTORC1 deficiency (Fig. 3C). Interestingly, reduced plasmablast frequency was found in the mLNs of immunized μMT:Cre^ER^*Rraga*^fl/fl^*Rragb*^fl/fl^ mice, but not in those of μMT:Cre^ER^*Rptor*^fl/fl^ mice (Fig. S3F), suggesting a potential tissue specific plasmablast defect in the absence of Raptor. Consistent with the significant reduction of GC and plasmablast generation, total and high affinity NP-specific antibodies of all classes were highly reduced in both knockout strains. Intriguingly, antibody titers from μMT:Cre^ER^*Rraga*^fl/fl^*Rragb*^fl/fl^ mice were consistently, although not significantly, lower than those from μMT:Cre^ER^*Rptor*^fl/fl^ mice (Fig. 3D-3E), which could be a reflection of the milder plasmablast defect induced by Raptor deletion relative to that induced by RagA/RagB deletion. Finally, we accessed p-S6 and p-4EBP1 in splenic B cells, either at steady state, or after overnight stimulation with anti-IgM/anti-CD40/IL-4, from immunized μMT:Cre^ER^*Rraga*^fl/fl^*Rragb*^fl/fl^ mice and μMT:Cre^ER^*Rptor*^fl/fl^ mice. We did not observe significant reduction of mTORC1 activity in RagA/RagB deficient B cells in either condition (Fig. 3F). Taken together, these data indicate an mTORC1 independent function of Rag-GTPases following immunization and illustrate a stronger dependence on Rag-GTPases than on mTORC1 for plasmablast generation.

**Figure 3.**
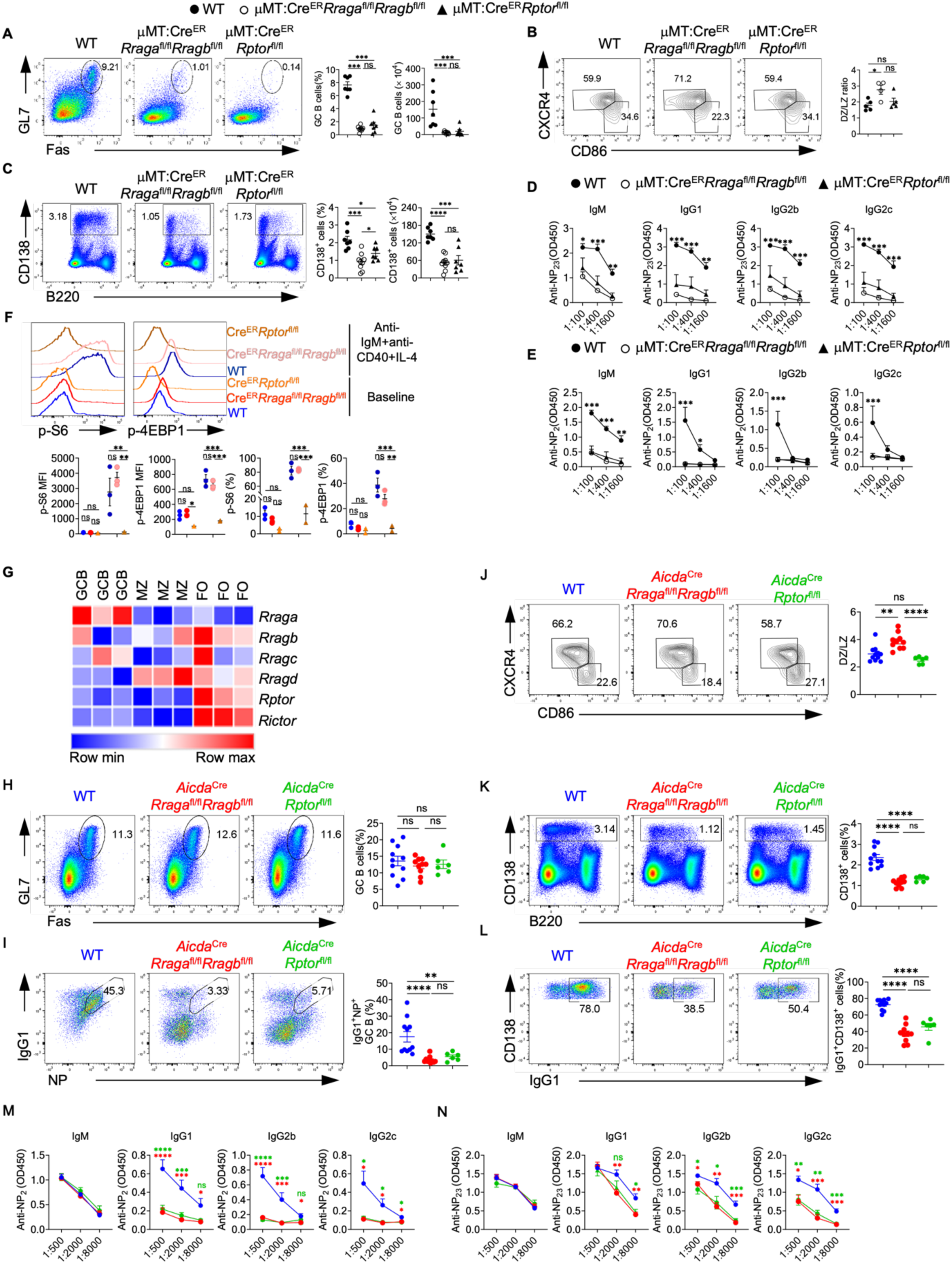
Rag-GTPases are critically required for GC formation and antibody production independent of mTORC1. (A-E) Tamoxifen was administered to animals by oral gavage daily for 4 consecutive days. Mice were immunized intraperitoneally (100 μg NP-OVA/alum) 7 days after the last injection. (A) Representative flow plots of GL-7 and Fas expression in splenic B cells from μMT:Cre^ER^ control (WT, n = 7), μMT:Cre^ER^ *Rraga*^fl/fl^ *Rragb*^fl/fl^ (n = 9), and μMT:Cre^ER^ *Rptor*^fl/fl^ (n = 7) mice. Right, summary of the percentages and numbers of GC (GL-7^+^ Fas^+^) B cells. (B) Representative flow plots of CXCR4 and CD86 expression in GC B cells. Right, summary of the ratio between dark zone (DZ) (CXCR4^+^ CD86^−^) and light zone (LZ) (CXCR4^−^ CD86^+^) cells. (C) Representative flow plots of CD138 and B220 expression in spleens of WT (n = 7), μMT:Cre^ER^ *Rraga*^fl/fl^ *Rragb*^fl/fl^ (n = 9), and μMT:Cre^ER^ *Rptor*^fl/fl^ (n = 7) mice. Right, summary of the percentages and numbers of CD138 cells. (D) Total NP-specific antibodies of all classes were measured using the sera of immunized WT (n = 7), μMT:Cre^ER^ *Rraga*^fl/fl^ *Rragb*^fl/fl^ (n = 8), and μMT:Cre^ER^ *Rptor*^fl/fl^ (n = 7) mice. (E) high-affinity NP-specific antibodies of all classes were measured using the sera of immunized WT (n = 5), μMT:Cre^ER^ *Rraga*^fl/fl^ *Rragb*^fl/fl^ (n = 5), and μMT:Cre^ER^ *Rptor*^fl/fl^ (n = 7) mice. (F) Representative flow plots of p-S6 and p-4EBP1 expression in fresh or overnight anti-IgM/anti-CD40/IL-4 stimulated splenic B cells from immunized WT (n = 3), μMT:Cre^ER^ *Rraga*^fl/fl^ *Rragb*^fl/fl^ (n = 3), and μMT:Cre^ER^ *Rptor*^fl/fl^ (n = 2) mice. (G) Heatmap of the mRNA expression of indicated genes on mouse follicular (FO) B cells, marginal zone (MZ) B cells and germinal center (GC) B cells. (H-N) NP-OVA was administered to WT (n = 11), *Aicda Rraga*^fl/fl^ *Rragb*^fl/fl^ (n = 10), and *Aicda Rptor*^fl/fl^ (n = 6) mice. Animals were analyzed 9 days after immunization. (H) Representative flow plots of GL-7 and Fas expression in splenic B cells. Right, summary of the GC B cell percentages. (I) Representative flow cytometry plots of NP^+^ IgG1^+^ GC B cells in splenic B cells. Right, summary of the percentages of NP^+^ IgG1^+^ GC B cells. (J) Representative flow plots of CXCR4 and CD86 expression in GC B cells. Right, summary of the DZ/LZ ratio. (K) Representative flow plots of CD138 and B220 expression in splenic lymphocytes. Right, summary of the percentages of CD138^+^ cells. (L) Representative flow plots of CD138 and IgG1^+^ expression in plasmablasts. Right, summary of the percentages of IgG1 plasmablasts. (M-N) The anti-NP_2_ (M) and anti-NP_23_ (N) titers of different immunoglobulin isotypes in the sera of immunized mice were measured by ELISA. Data in graphs represent mean ± SEM. ns, not significant. *p < 0.05, **p < 0.01, ***p < 0.001, and ****p < 0.0001, one-way ANOVA (A, B, C, H, I, J, K and L), two-way ANOVA (D, E, F, M and N).

The above defective GC formation in the absence of RagA/RagB or Raptor could be due to impaired B cell activation. Moreover, we found that *Rraga*, among all Rag family members and mTOR scaffolding molecules, preferentially expressed in GC B cells (Fig. 3G), suggesting a prominent role of RagA in the GC response. To investigate the role of Rag-GTPases post B cell activation, we generated *Aicda*^Cre^*Rraga*^fl/fl^*Rragb*^fl/fl^ mice and *Aicda*^Cre^*Rptor*^fl/fl^ mice to ablate RagA/RagB and Raptor after B cell activation, especially in GC B cells(*54*). After immunization with NP-OVA/alum, we did not find any apparent alteration of the GC B cell frequencies in the spleens of either *Aicda*^Cre^*Rraga*^fl/fl^*Rragb*^fl/fl^ mice or *Aicda*^Cre^*Rptor*^fl/fl^ mice compared with WT mice (Fig. 3H). However, NP^+^IgG1^+^ GC B cells highly reduced in the spleens of both mouse strains (Fig. 3I), indicating that Rag-GTPases and mTORC1 are both critical for antigen selection in GC B cells. Furthermore, immunized *Aicda*^Cre^*Rraga*^fl/fl^*Rragb*^fl/fl^ mice harbored higher DZ/LZ ratio, while a slightly reduced DZ/LZ ratio was observed in immunized *Aicda*^Cre^*Rptor*^fl/fl^ mice (Fig. 3J). We also observed substantially reduced CD138^+^ plasmablast frequency and IgG1 expression on plasmablasts in both *Aicda*^Cre^*Rraga*^fl/fl^*Rragb*^fl/fl^ and *Aicda*^Cre^*Rptor*^fl/fl^ mice (Fig. 3K-3L).

Measurement of NP-specific antibodies showed decreased titers of high-affinity and total IgG isotypes, but not IgM, from the sera of either mouse strain (Fig. 3M-3N). Altogether, these data demonstrate that both Rag-GTPases and mTORC1 are required for TD antigen induced GC formation. But Rag-GTPases and mTORC1 likely employ distinct mechanisms to promote plasmablast formation and maintain GC dynamics.

### Rag-GTPases support mitochondrial fitness during B cell activation, independent of mTORC1

B cell activation is accompanied by extensive metabolic reprogramming including glycolytic switches and activation of mitochondrial oxidative phosphorylation (*55*). We examined mitochondrial respiration and glycolysis in activated B cells by measuring oxygen consumption rate (OCR) and extracellular acidification rate (ECAR), respectively. Deficiency of either RagA/RagB or Raptor led to significantly decreased OCR (Fig. 4A). Raptor deficiency strongly suppressed glycolysis, while RagA/RagB deficient B cells had a slight reduction of ECAR (Fig. 4B). [3-^3^H]-glucose labeling assay confirmed the profound glycolytic defect in Raptor deficient B cells. It showed a modest but significant reduction of glucose metabolism in RagA/RagB deficient B cells (Fig. 4C). Thus, Rag-GTPases are critical for B cell metabolism, especially oxidative phosphorylation. Interestingly, despite a relatively stronger reduction of OCR in Raptor deficient B cells, RagA/RagB deficiency, but not Raptor deficiency, resulted in significant reduction of mitochondrial membrane potential, measured by tetramethylrhodamine methyl ester (TMRM) (Fig. 4D) and MitoTracker Deep Red (MTDR) (Fig. 4E) (*56, 57*), as well as mitochondrial reactive oxygen species (ROS) measured by MitoSox (Fig. 4F), suggesting that loss of Rag-GTPases might lead to defective mitochondria and subsequent reduced mitochondrial activity, while mTORC1 deficiency compromises OCR through distinct mechanisms. Moreover, examination of mitochondrial phenotypes *in vivo* revealed that GC B cells from the immunized *Aicda*^Cre^*Rraga*^fl/fl^*Rragb*^fl/fl^ mice, but not *Aicda*^Cre^*Rptor*^fl/fl^ mice, showed significantly reduced mitochondrial membrane potential and ROS level (Fig. 4G-4H). Previous studies indicated that mTORC1 controlled the expression of key transcription factors for mitochondrial biogenesis program, including PGC-1α and TFAM (*58, 59*). Indeed, expression of PGC-1α (Fig. 4I), TFAM (Fig. 4J) (*60*), and COXIV (Fig. 4K), a member of the mitochondrial respiratory chain were all significantly decreased in Raptor deficient B cells, but not in RagA/RagB deficient B cells. Collectively, Rag-GTPase deficiency in B cells impairs mitochondrial metabolism associated with reduced mitochondrial membrane potential and mitochondrial ROS, which are distinctive from Raptor deficient B cells.

**Figure 4.**
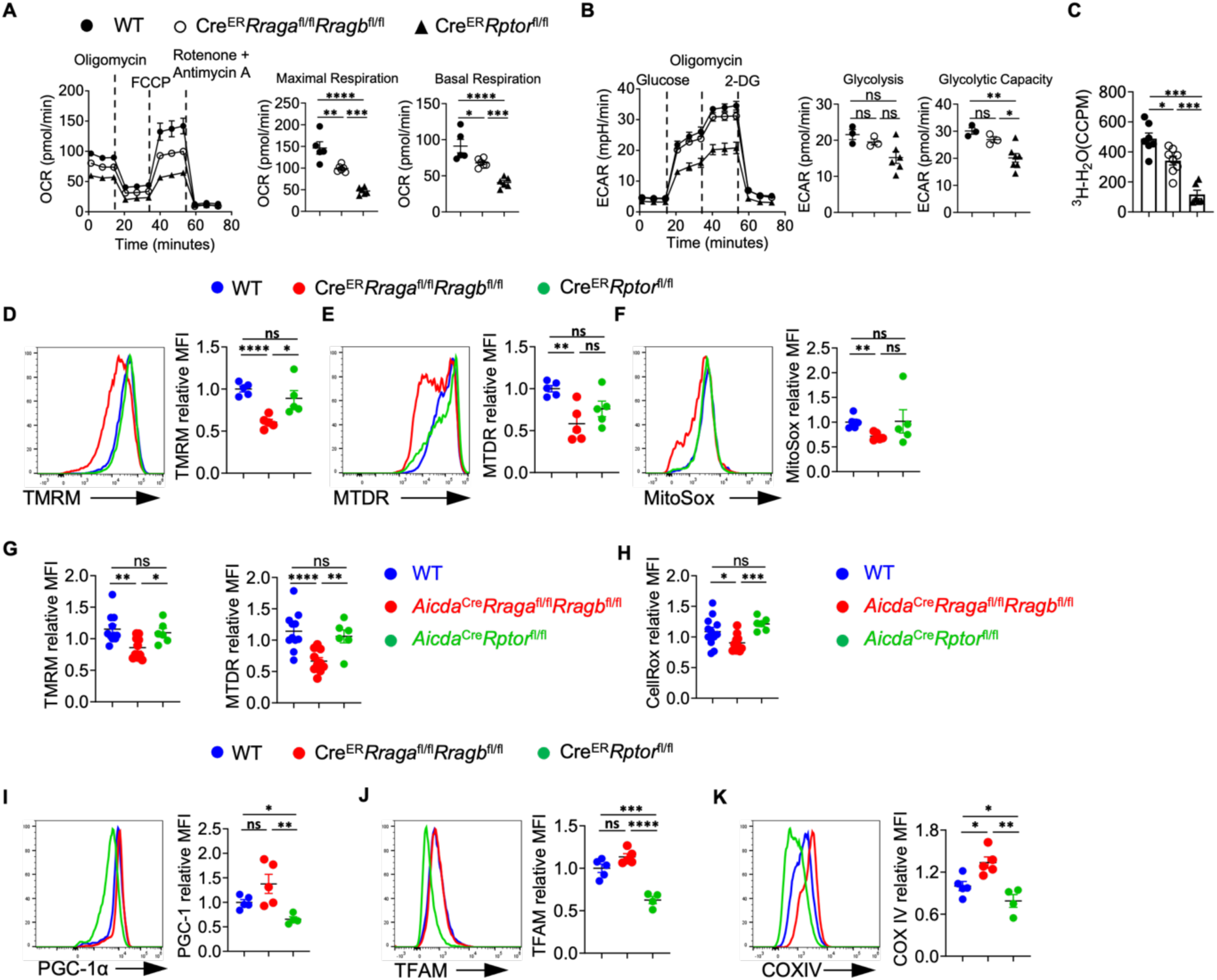
Rag-GTPases regulate B cell activation and mitochondrial metabolism differently from canonical mTORC1. (A-F, I-K) Splenic B cells were purified from tamoxifen-treated WT, *Cre*^ER^ *Rraga*^fl/fl^ *Rragb*^fl/fl^, and *Cre*^ER^ *Rptor*^fl/fl^ mice, and cultured with LPS/IL-4/BAFF for 72 h. Mitostress assay (A) and glycolytic stress assay (B) were performed on a Seahorse XFe96 bioanalyzer. (C) Glycolytic flux was examined using LPS/IL-4/BAFF activated B cells by measuring the detritiation of [3-H] glucose. Representative flow plots of tetramethylrhodamine methyl ester (TMRM) (D, 4-5 mice per group), MitoTracker Deep Red (MTDR) (E, 4-5 mice per group), and MitoSox (F, 4-5 mice per group) stainings were shown. Summaries of the mean fluorescence intensity (MFI) of each staining (relative to WT) were on the right. (G) Summaries of the MFIs of TMRM and MTDR staining (relative to WT) on GC B cells from the spleens of immunized mice. (H) Summay of the MFIs of CellRox staining (relative to WT) on GC B cells from the spleens of immunized. Representative flow plots of PGC-1α (I, 4-5 mice per group), TFAM (J, 4-5 mice per group) and COXIV (K, 4-5 mice per group) stainings were shown. Summaries of the MFI of each staining (relative to WT) were on the right. Data in graphs represent mean ± SEM. ns, not significant. *p < 0.05, **p < 0.01, ***p < 0.001, and ****p < 0.0001, one-way ANOVA (A-K).

### Rag-GTPase deficiency leads to TFEB/TFE3 overactivation and abnormal mitophagy

To probe the molecular mechanisms underlying the above metabolic defects, we conducted RNA sequencing using activated B cells and GC B cells. Gene set enrichment analysis (GSEA) identified the KEGG lysosome pathway and putative TFEB target genes as two of the top enriched pathways in RagA/RagB deficient B cells (Fig. 5A-5B, Fig. S4A). Many lysosomal genes were upregulated in the RagA/RagB deficient B cells (Fig. S4B), such as *Lamp1*, *Atp6ap1*, *Atp6v1h* and *Ctsa*. RNA sequencing using GC B cells from the immunized *Aicda*^Cre^*Rraga*^fl/fl^*Rragb*^fl/fl^ mice and *Aicda*^Cre^*Rptor*^fl/fl^ mice demonstrated a relatively effective deletion of *Rraga* and *Rptor* (Fig. S4C). The PCA plot showed that WT and Raptor deficient cells were closer to each other, while both were distant from Rag-GTPases deficient cells (Fig. S4D). Consistent with this observation, Venn diagram of DEGs illustrated that majority of DEGs were from the RagA/RagB KO vs WT comparison (85.7%, 1812/2114), and less than 10% of DEGs (175/2114) were identified comparing Raptor KO with WT (Fig. 5C, Fig. S4E). There were more DEGs shared between RagA/RagB KO vs WT and RagA/RagB KO vs Raptor KO than between Raptor KO vs WT, indicating a stronger impact of Rag-GTPase deficiency on GC B cell transcriptome than Raptor deficiency. GSEA identified KEGG lysosome pathway as the top enriched pathway comparing RagA/RagB KO to WT GC B cells (Fig. 5D), or comparing RagA/RagB KO to Raptor KO GC B cells (Fig. 5E), while no significant enrichment of KEGG lysosome was observed when comparing GC B cells from *Aicda*^Cre^*Rptor*^fl/fl^ and WT mice (Fig. 5F), suggesting lysosomal activation in RagA/RagB deficient, but not in Raptor deficient, GC B cells. Therefore, Rag-GTPases and mTORC1 utilize distinct mechanisms to maintain GC reaction post B cell activation, one of which was likely TFEB regulation.

**Figure 5.**
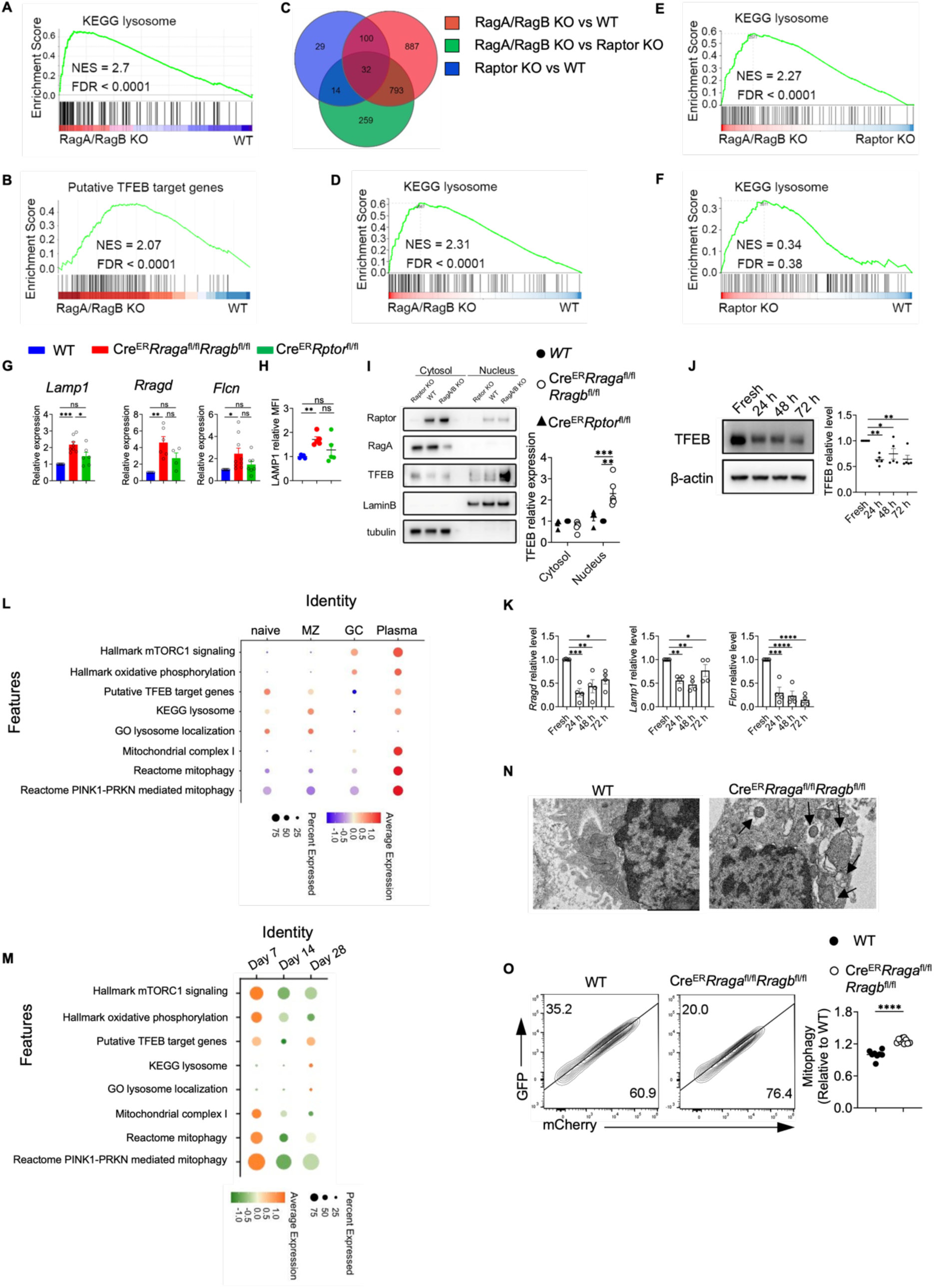
RagA/RagB deficiency leads to TFEB/TFE3 overactivation and abnormal mitophagy. (A) Bulk RNA sequencing was performed on the B cells activated with LPS/IL-4/BAFF for 72 h. Gene set enrichment analysis (GSEA) was conducted using differentially expressed genes (DEGs), and KEGG lysosome pathway was plotted. (B) The enrichment of the putative TFEB target genes was evaluated using the DEGs. (C) Bulk RNA sequencing was performed on the sorted GC B cells from the indicated genotypes. DEGs between different comparisons were calculated and the Venn diagram of three comparisons was presented. (D-F) GSEA was performed using the DEGs between indicated genotypes. KEGG lysosome pathway enrichement between RagA/RagB deficient and WT B cells (D), between RagA/RagB deficient and Raptor deficient B cells (E), and between Raptor deficient and WT B cells (F) were presented. (G) Splenic B cells were stimulated with LPS/IL-4/BAFF for 72h, then qRT-PCR was used to measure the expression of TFEB target genes *Lamp1*, *Rragd*, and *Flcn,* n = 4-9. (H) LAMP1 expression was measured by flow cytometry, n = 4-5 per group. (I) Cytosolic and nuclear proteins were isolated from the B cells activated with LPS/IL-4/BAFF for 72 h. Expression of Raptor, RagA and TFEB was examined by immunoblot. Lamin B was used as a nuclear control, tubulin was used as a cytosol control. (J) Expression of TFEB was examined by immunoblot using B cells activated with LPS/IL-4/BAFF for the indicated time (n = 5). Right, summary of TFEB expression (relative to fresh). (K) Expression of TFEB target genes *Lamp1*, *Rragd*, and *Flcn* was examined by qRT-PCR using B cells activated as in J. (L-M) scRNA seq data from the published database (E-MTAB-9478) was reanalyzed, and different gene signatures were evaluated on naïve B cells, marginal zone (MZ) B cells, germinal center (GC) B cells, and plasma cells (L). Expression of the indicated gene signatures on GC B cells at different time points (M). (N) Activated B cells were examined by transmission electron microscopy (TEM). Arrows indicate mitochondria surrounded by double layer structures. Scale bar = 2 μm. (O) Representative flow plots of GFP and mCherry expression in activated B cells transduced with Mito-QC. Right, summary of the normalized mCherry percentage (Relative to mCherry percentage in WT). Data in graphs represent mean ± SEM. ns, not significant. *p < 0.05, **p < 0.01, ***p < 0.001, and ****p < 0.0001, one-way ANOVA (G, H, I, J and K), two-tailed Student’s t test (O).

To directly examine TFEB function activity, we measured the mRNA levels of TFEB target genes, including *Lamp1*, *Rragd*, and *Flcn.* Consistent with the RNA sequencing data, TFEB target gene expression was significantly increased in RagA/RagB deficient B cells, but not in Raptor deficient B cells (Fig. 5G). We also confirmed that LAMP1 protein expression was increased in the absence of Rag-GTPases, but not Raptor (Fig. 5H, Fig. 1D). Importantly, we observed increased TFEB and TFE3 nuclear localization in RagA/RagB deficient B cells, but not in Raptor deficient B cells (Fig. 5I, Fig. S4F). Therefore, Rag-GTPases, but not mTORC1, constrain TFEB/TFE3 activity during B cell activation.

To further gain insight into the possible functions of TFEB during B cell activation, we first examined TFEB protein expression kinetics in *in vitro* activated B cells. TFEB protein had the highest expression in naïve B cells and its expression declined overtime (Fig. 5J). Meanwhile, we observed reduced expression of *Lamp1, Rragd*, and *Flcn* (Fig. 5K), indicating an association between B cell activation and reduced TFEB activity. To further investigate the TFEB activity *in vivo*, we re-analyzed the public database and evaluated the expression of mTORC1 signaling signatures, putative TFEB target genes (*61*), oxidative phosphorylation, and lysosome related genes in naïve B cells, MZ B cells, GC B cells and plasma cells (Fig. 5L) and GC B cells at different time points (Fig. 5M) from influenza infected mice. While mTORC1 signatures, oxidative phosphorylation, and mitochondrial complex I were highly enriched in plasma cells, TFEB targets and lysosome related genes were enriched in naïve and MZ B cells, and they had the lowest expression in GC B cells (Fig. 5L). In the time-course analysis, TFEB target gene expression declined at day 14 compared to day 7 before recovering at day 28 (Fig. 5M). The expression pattern and kinetics of TFEB activity were not reciprocal to mTORC1 activation, suggesting that they could be independent to each other in B cells. Taken together, our results suggest that B cell activation and differentiation are associated with the decline of TFEB activity, which might be enforced by Rag-GTPases.

TFEB has been implicated in the regulation of mitophagy (*62, 63*). Yet, the function of TFEB and mitophagy during B cell activation remains unknown. Transmission electron microscopy (TEM) analysis revealed mitochondria encircled with double membrane structures, morphology consistent with mitophagy, in RagA/RagB deficient B cells (Fig. 5N). To confirm the mitophagy phenotype, we introduced Mito-QC, a mitophagy reporter, which is a construct expressing an mCherry-GFP tag attached to mitochondrial fission protein 1 (FIS1, residues 101–152) on the outer mitochondrial membrane (*64*). Upon mitophagy, mitochondria are delivered to lysosomes from autophagosomes, where the GFP signal is quenched by the low lysosomal pH, resulting in only red fluorescence (mCherry signals). In accordance with the TEM data, we observed increased mCherry signals in RagA/RagB deficient B cells (Fig. 5O). Thus, Rag-GTPase deficiency is associated with increased TFEB activity and mitophagy in B cells.

### TFEB/TFE3 overactivation is responsible for the abnormal mitophagy and impaired mitochondrial fitness in RagA/RagB deficient B cells

To test whether TFEB overactivation can directly control mitochondrial phenotypes and B cell activation, we retrovirally overexpressed WT TFEB or constitutive active TFEB (Ca TFEB) carrying S142/211A mutation (Fig. S5A) (*65*), with comparable transduction efficiency (Fig. S5B). WT TFEB and Ca TFEB overexpression promoted TFEB activation in a graded manner, as measured by the expression of TFEB target genes (Fig. 6A). They also reduced IgG1^+^ class switching (Fig. 6B) and CD138^+^ expression (Fig. S5C) in a graded manner, without affecting B cell proliferation (Fig. S5D). We observed similar phenotypes when we pharmacologically stimulated TFEB using curcumin analog C1, a novel mTOR-independent activator of TFEB (*66*) (Fig. S5E-S5F). Hence, TFEB overactivation suppresses class switch and CD138 expression *in vitro*.

**Figure 6.**
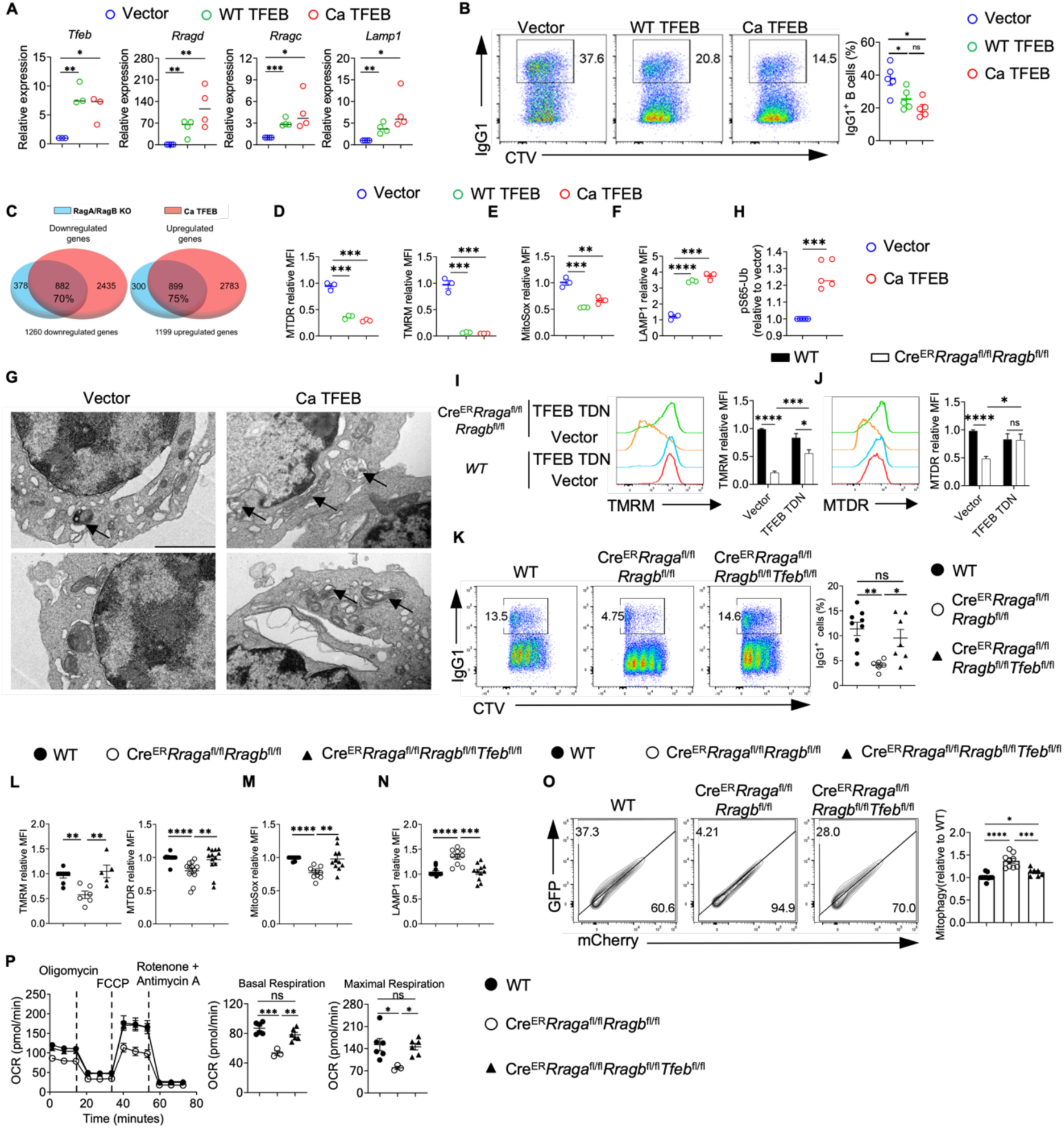
TFEB/TFE3 overactivation is responsible for the abnormal mitophagy and impaired mitochondrial fitness in RagA/RagB deficient B cells. (A) GFP B cells were sorted from the vector, WT TFEB, and Ca TFEB transduced B cells. Expression of the TFEB target genes was examined by qRT-PCR. (B) Representative flow plots of IgG1 expression and Celltrace violet (CTV) dilution in vector, WT TFEB, or Ca TFEB transduced GFP^+^ B cells. (C) Venn diagrams highlight the overlapping gene numbers in downregulated DEGs (left) and upregulated DEGs (right) between Rag KO vs WT comparison and Ca TFEB vs vector analysis. (D) Summaries of the relative MTDR (left) and TMRM (right) staining MFIs. (E) Summary of the relative MitoSox staining MFIs. (F) Summary of the relative LAMP1 staining MFIs. (G) GFP B cells from vector or Ca TFEB transduced B cells were sorted and subjected to transmission electron microscope (TEM) analysis. Arrows indicate mitochondria surrounded by double layer structures., scale bar = 2 μm. (H) GFP B cells from vector or Ca TFEB transduced B cells were sorted, and ELISA was conducted to measure the level of pS65-Ub. (I-J) B cells from WT (n = 5) and *Cre*^ER^ *Rraga*^fl/fl^ *Rragb*^fl/fl^ (n = 7) mice were transduced with vector or TFEB TDN. Representative flow plots of TMRM (I) or MTDR (J) staining were presented. (K) B cells were labeled with CTV and activated with LPS/IL-4/BAFF for 3 days, expression IgG1 and CTV dilution were examined by flow cytometry. (L-N) B cells from indicated genotypes were activated with LPS/IL-4/BAFF for 3 days, TMRM or MTDR (L), MitoSox (M) or LAMP1 (N) were measured by flow cytometry. (O) Mito-QC system was used on the indicated cells to check the mitophagy. *Cre*^ER^ *Rraga*^fl/fl^ *Rragb*^fl/fl^ (n = 9), *Cre*^ER^ *Rraga*^fl/fl^ *Rragb*^fl/fl^ Tfeb^fl/fl^ (n = 7), and WT (n = 10). (P) B cells from the indicated mice were purified and activated with LPS/IL-4/BAFF for 72 h. Mitostress assay was performed on a Seahorse XFe96 analyzer. Right, summaries of the basal respiration and maximal respiration. Data in graphs represent mean ± SEM. ns, not significant. *p < 0.05, **p < 0.01, ***p < 0.001, and ****p < 0.0001, one-way ANOVA (A, B, D, E, F, K, L, M, N, O and P), two-way ANOVA (I and J), two-tailed Student’s t test (H).

RNA sequencing revealed the enrichment of lysosome pathway in Ca TFEB transduced B cells (Fig. S5G). There was a substantial overlap between the differential expressed genes (DEGs) from the RagA/RagB deficient B cells and those from Ca TFEB expressing B cells: 70% downregulated genes (882 genes) and 75% upregulated genes (899 genes) were shared (Fig. 6C), suggesting that TFEB overactivation might account for a significant portion of the DEGs in RagA/RagB deficient B cells. Like RagA/RagB deficient B cells, TFEB overactivation led to striking reductions of mitochondrial membrane potential (TMRM and MTDR, Fig. 6D) and mitochondrial ROS (MitoSox, Fig. 6E), and increased LAMP1 expression (Fig. 6F). Importantly, TEM analysis revealed the increased mitochondrial morphology consistent with mitophagy in Ca TFEB transduced B cells (Fig. 6G). Of note, TFEB overactivation promoted mitophagy likely through PINK1-PRKN/Parkin mediated pathway because we detected a significant increase of ubiquitin (Ub) phosphorylation at Ser65 (p-S65-Ub)(*67*) (Fig. 6H). Thus, TFEB activation is sufficient to suppress mitochondrial activity associated with abnormal mitophagy in B cells.

To establish whether TFEB overactivation was responsible for B cell activation and mitochondrial defects observed in RagA/RagB deficient B cells, we generated dominant negative TFEB (TFEB TDN), which contained the helix-loop-helix–leucine zipper dimerization domains but lacked the DNA-binding basic region and transcription activation domains (*68*). TFEB TDN transduction partially suppressed TFEB overactivation and partially rescued the reduced IgG1^+^ expression in RagA/RagB deficient B cells (Fig. S5H-S5I). Moreover, TFEB TDN partially restored mitochondrial membrane potential (Fig. 6I-6J) and MitoSox level in RagA/RagB deficient B cells (Fig. S5J). These data indicate that TFEB overactivation could be partly responsible for the mitochondrial defects induced by RagA/RagB deficiency *in vitro*.

To investigate if genetic inactivation of TFEB can restore B cell mitochondrial fitness in the absence of Rag-GTPases, we crossed Cre^ER^*Rraga*^fl/fl^*Rragb*^fl/fl^ mice with a *Tfeb*^fl/fl^ allele to generate Cre^ER^*Rraga*^fl/fl^*Rragb*^fl/fl^*Tfeb*^fl/fl^ mice. As expected, loss of TFEB greatly reduced the expression of *Tfeb* and many TEFB target genes in RagA/RagB deficient B cells (Fig. S5K). TFEB deletion restored the reduced IgG1 class switch (Fig. 6K), mitochondrial membrane potential (Fig. 6L), mitochondrial ROS (Fig. 6M), and reversed the increased expression of LAMP1 (Fig. 6N). Importantly, TFEB deficiency prevented the abnormal mitophagy measured by Mito-QC assay (Fig. 6O). Finally, the reduced mitochondrial metabolism, measured by OCR, induced by RagA/RagB loss was fully restored by TFEB deletion (Fig. 6P). Altogether, these data demonstrated that Rag-GTPases constrain TFEB/TFE3 activity to prevent abnormal mitophagy and maintain mitochondrial fitness in B cells *in vitro*.

### Rag-GTPase-TFEB/TFE3 axis modulates humoral immunity in a context dependent manner

We next sought to assess the impact of TFEB deletion on Cre^ER^*Rraga*^fl/fl^*Rragb*^fl/fl^ mice *in vivo*. TFEB deletion restored the reduced frequencies of fractions B and fraction C/C’ precursors, but not the blockage of the pro-B to pre-B transition, nor the reduced GC formation in PPs caused by RagA/RagB deficiency (Fig. S6A-S6C). To evaluate the humoral immune responses against TD antigens, we immunized tamoxifen injected μMT:Cre^ER^*Rraga*^fl/fl^*Rragb*^fl/fl^*Tfeb*^fl/fl^ mice and μMT:Cre^ER^*Rraga*^fl/fl^*Rragb*^fl/fl^ mice with NP-OVA/alum. TFEB deletion did not restore the reduced GC B cells, plasmablasts, NP^+^IgG1^+^ GC B cells, IgG1^+^CD138^+^ cells and the production of NP specific antibodies in the absence of RagA/RagB (Fig. S6D-S6I). Thus, TFEB deletion was not sufficient to restore early B cell development, GC formation in PPs and humoral responses towards TD antigens in the absence of Rag-GTPases.

To investigate immune response to TI-antigen, we immunized tamoxifen injected μMT:Cre^ER^*Rraga*^fl/fl^*Rragb*^fl/fl^*Tfeb*^fl/fl^ mice and μMT:Cre^ER^*Rraga*^fl/fl^*Rragb*^fl/fl^ mice with TNP-LPS. RagA/RagB deletion led to significant reduction of plasmablast generation and TNP specific antibody production, both of which were restored by TFEB deletion (Fig. S6J-S6K). Interestingly, we observed that plasmablasts generated by TNP-LPS immunization exhibited greater mitochondrial membrane potential, measured by TMRM and MTDR, than those generated by NP-OVA immunization (Fig. S6L), suggesting a potentially higher mitochondrial metabolism in TNP-LPS induced plasmablasts than NP-OVA induced plasmablasts. Thus, our data indicate that TFEB overactivation is responsible for the impaired TI-antigen responses, but not the reduced TD-antigen responses, pro-B to pre-B transition, or GC formation in PPs in the absence of Rag-GTPases.

RagA/RagB deficiency led to overactivation of both TFEB and TFE3 (Fig. 5I, Fig. S4F). To address the potential redundancy between TFEB and TFE3, we generated Cre^ER^*Rraga*^fl/fl^*Rragb*^fl/fl^*Tfeb*^fl/fl^*Tfe3*^−/−^ mice. Strikingly, we found that TFEB/TFE3 deletion significantly restored the reduction of B220^lo^CD43^−^ pre-B/immature B cells and fraction B and fraction C/C’ frequencies in the BM (Fig. 7A-7B), and GC formation in PPs (Fig. 7C) in the absence of RagA/RagB, demonstrating a non-redundant role between TFE3 and TFEB during early B cell development and spontaneous GC formation in mucosal site under the control of Rag-GTPases. Like TFEB deletion, TFEB/TFE3 deletion restored most of the *in vitro* activation and metabolic defects in RagA/RagB deficient B cells, including reduced IgG1 expression (Fig. 7D), mitochondrial membrane potential (Fig. 7E), ROS level (Fig. 7F) and increased LAMP1 expression (Fig. 7G), as well as the reduced OCR (Fig. 7H). Next, we immunized tamoxifen injected μMT:Cre^ER^*Rraga*^fl/fl^*Rragb*^fl/fl^*Tfeb*^fl/fl^*Tfe3*^−/−^ mice and μMT:Cre^ER^*Rraga*^fl/fl^*Rragb*^fl/fl^ mice with NP-OVA/alum or TNP-LPS. TFEB/TFE3 deficiency was not able to rescue the reduced GC and plasmablast formation (Fig. 7I-7J) upon NP-OVA immunization in the absence of RagA/RagB. However, the reduced responses towards TNP-LPS immunization in RagA/RagB deficient B cells were nearly restored by TFEB/TFE3 deletion (Fig. 7K-7L). Taken together, these data reveal non-redundant functions for TFEB and TFE3 in B cell development and GC formation in mucosal site. They support the notion that Rag-GTPase-TFEB/TFE3 axis regulates B cell development, activation, and differentiation under different immune context.

**Figure 7.**
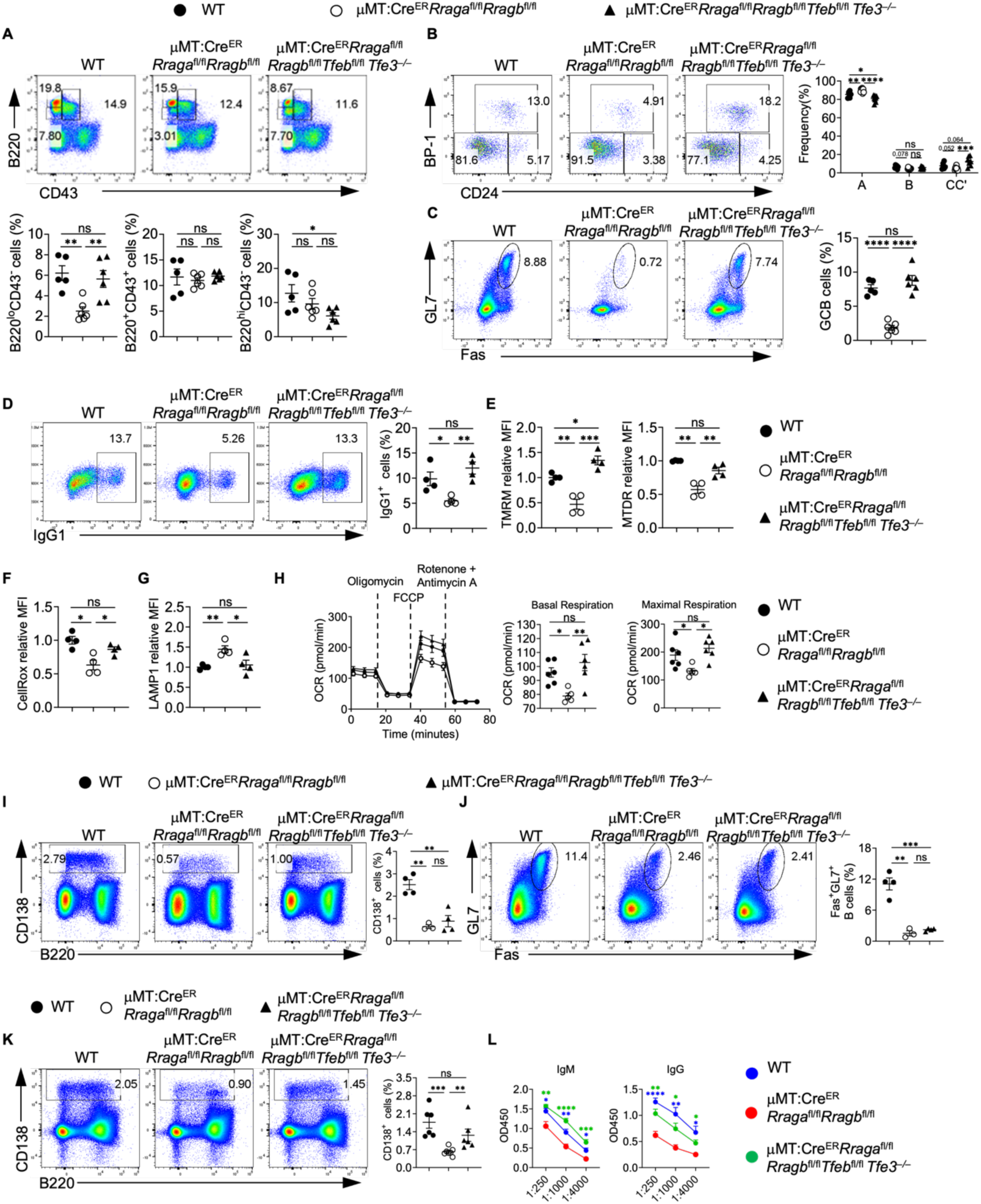
Rag-GTPases-TFEB/TFE3 axis modulates humoral immunity in a context dependent manner. (A) Representative flow plots of CD43 and B220 expression on BM cells. Right, summaries of the frequencies of B220^lo^ CD43^−^, B220^hl^ CD43^−^ and B220^+^ CD43^+^ cells. (B) Representative flow plots of BP-1 and CD24 expression in BM B220^+^ CD43^+^ IgM^−^ B cell precursors. Right, summary of the frequencies of fraction A (CD24^−^ BP-1^−^), fraction B (CD24^+^ BP-1^+^), and fraction C/C′ (CD24 BP-1) cells. (C) Representative flow plots of GL-7 and Fas expression in lymphocytes from the Peyer’s patches. Right, summary of the frequencies of GC B cells. (D-E) Splenic B cells from the indicated chimera mice were purified and stimulated with LPS/IL-4/BAFF for 72 h. IgG1 expression (D), TMRM and MTDR staining (E), CellROX staining (F) and LAMP1 staining (G) were examined by flow cytometry. Summaries of the frequencies of IgG1 B cells, the relative TMRM, MTDR, CellROX, and LAMP1 staining MFIs were presented. (H) Mitostress assay was performed on a Seahorse XFe96 analyzer. Right, summaries of the basal respiration and maximal respiration. (I-L) Tamoxifen injected WT, μMT:Cre^ER^ *Rraga Rragb*^fl/fl^, and μMT:Cre^ER^ *Rraga*^fl/fl^ *Rragb*^fl/fl^ *Tfeb*^fl/fl^ *Tfe3^-/-^* chimera mice were immunized intraperitoneally with 100 μg NP-OVA/alum (I-J) or 50 μg TNP-LPS (K-L). (I) Representative flow plots of CD138^+^ and B220 expression in splenic lymphocytes. Right, summary of the frequency of CD138 plasmablasts. (J) Representative flow plots of GL-7 and Fas expression on splenic B cells. Right, summary of the frequency of GC B cells. (K) Representative flow plots of CD138 and B220 expression in splenic lymphocytes. Right, summary of the frequency of CD138 plasmablasts. (L)Titers of TNP-specific antibodies in the sera of immunized mice were measured by ELISA. Data in graphs represent mean ± SEM. ns, not significant. *p < 0.05, **p < 0.01, ***p < 0.001, and ****p < 0.0001, one-way ANOVA (A, C, D, E, F, G, H, I, J and K), two-way ANOVA (B and L).

## Discussion

Research in childhood malnutrition has long highlighted the impact of nutrient availability and immunity. Yet, the complex interplay between systemic metabolic dysregulation and immune system hampers the mechanistic understanding of nutrient sensing and immunometabolism in adaptive immune system (*69*). Recent studies have established Rag-GTPases as a key sensor for amino acids. Rag-GTPases primarily control mTORC1 and TFEB/TFE3 activity. In the current study, we utilized genetic models to establish the mechanisms through which Rag-GTPases suppress TFEB/TFE3 in B cell development and activation in an mTORC1 independent manner.

The relationship between Rag-GTPases and mTORC1 has been contentious. Rag-GTPases were initially found to be necessary and sufficient for mTORC1 activation, including in immune cells (*21, 23, 24, 32*). However, amino acids can engage mTORC1 independent of Rag-GTPases(*30*). Our study demonstrated that Rag-GTPases are not necessary for mTORC1 activation in B cells. Our observations join several recent studies to illustrate the relationship between Rag-GTPases and mTORC1 is cell type and context dependent: Rag-GTPases can be sufficient but not necessary for mTORC1 activation in B cells (*24, 25*). Furthermore, Rag-GTPases and mTORC1 in GC B cells are critical for antigen selection and antibody affinity maturation. Yet only Rag-GTPases, but not mTORC1, modulate DZ vs LZ distribution and lysosome metabolism in GC B cells. Finally, Rag-GTPases play a more prominent role in supporting plasmablast formation than mTORC1, whose underlying mechanisms remain to be investigated. These results uncover the distinct requirements of Rag-GTPases and mTORC1 in humoral immunity.

While recent advances in immunometabolism field have identified certain metabolic requirements during mature B cell activation, much less is known about metabolic regulations during B cell development, spontaneous GC reaction in mucosal site and humoral responses towards TD and TI antigens. Our data establish a novel Rag-GTPase-TFEB/TFE3-mitophagy pathway that controls mitochondrial fitness in B cells. Instead of promoting mTORC1, Rag-GTPases suppress TFEB/TFE3 activity following B cell activation. Deficiency of Rag-GTPases leads to TFEB/TFE3 nuclear accumulation and overactivation, which induces mitophagy and reduces mitochondrial membrane potential and mitochondrial oxidative phosphorylation. Importantly, our data demonstrate a key differential requirement of TFEB/TFE3 transcription factors between TD and TI responses, i.e., Rag-GTPase mediated TFEB/TFE3 suppression is critical for TI response, but a Rag-GTPase dependent but TFEB/TFE3 independent mechanism is required for TD response. The molecular mechanisms governing TI responses are much less understood compared to those regulating TD responses. A recent study showed that TD, but not TI, responses depend on LDHA mediated glycolysis (*10*). Here, our study demonstrated that mitochondrial integrity mediated by Rag-GTPase-TFEB/TFE3 axis is critical for TI response but may not be sufficient for TD response. The TFEB/TFE3 independent but Rag-GTPase dependent mechanisms for TD response awaits future study. The higher mitochondrial membrane potential in plasmablasts induced by TI-antigen than TD-antigen suggests a possibility that TI-response might have a greater reliance on mitochondrial metabolism than TD-response, consistent with a recent study (*70*). More research is warranted to test this proposition. Thus, our investigation help fill a critical knowledge gap and further highlight the distinct signaling and metabolic requirements between TD and TI responses.

Finally, our study unveils overlapping and non-redundant functions between TFEB and TFE3 in B cells. While deletion of TFEB alone or both TFEB and TFE3 can restore the impaired antibody response to TI antigens in RagA/RagB deficient B cells, deletion of both TFEB and TFE3 is needed to rectify early B cell development defects and reduced GC formation in PPs in the absence of Rag-GTPases. Altogether, our investigations illustrate specific metabolic and signaling requirements in the lifetime of B cells at different stages and different anatomic locations coordinated by Rag-GTPase-TFEB/TFE3 signaling axis.

## Material and methods

### Mice

*ROSA26-*Cre-ERT2, *Rraga*^fl/fl^*Rragb*^fl/fl^, *Rptor^fl/fl^* mice have been previously described(*27, 34*). CD45.1^+^ (RRID: IMSR_JAX:002014), C57BL/6J (RRID: IMSR_JAX:000664), and *Rag1*^−/−^ (RRID: IMSR_JAX:002216), B6.129S2-*Ighm^tm1Cgn^*/J (RRID:IMSR_JAX:002288) and B6.129P2-*Aicda^tm1(cre)Mnz^*/J (RRID:IMSR_JAX:007770) mice were purchased from the Jackson Laboratory. *Tfeb*^fl/fl^ mouse was a gift from Dr. Andrea Ballabio (Telethon Institute of Genetics and Medicine)(*71*). *Tfe3^−/−^*mouse was a gift from Dr. Ming O. Li (Memorial Sloan Kettering Cancer Center)(*31*). *Cre*^ER^*Rraga*^fl/fl^*Rragb*^fl/fl^ (*Rragb*^fl/fl^ denotes male hemizygous or female homozygous mice for *Rragb* because *Rragb* is located on the X-chromosome). *Cre*^ER^*Rragb*^fl/fl^ mice, *Cre*^ER^*Rptor*^fl/fl^, *Cre*^ER^*Rraga*^fl/fl^*Rragb*^fl/fl^*Tfeb*^fl/fl^, *Cre*^ER^*Rraga*^fl/fl^*Rragb*^fl/fl^*Tfeb*^fl/fl^*Tfe3^−/−^*and age-and gender-matched littermate controls were analyzed at the indicated ages. Bone marrow (BM) chimeras were generated by transferring T cell-depleted bone marrow cells into lethally irradiated (11 Gy) CD45.1^+^ mice by mixing Cre^ER^*Rraga*^+/+^*Rragb*^+/+^*Rptor*^+/+^ (WT), *Cre*^ER^*Rraga*^fl/fl^*Rragb*^fl/fl^, *Cre*^ER^*Rraga*^fl/fl^*Rragb*^fl/fl^*Tfeb*^fl/fl^, *Cre*^ER^*Rraga*^fl/fl^*Rragb*^fl/fl^*Tfeb*^fl/fl^*Tfe3^−/−^* or *Cre*^ER^*Rptor*^fl/fl^ BM with μMT BM at a ratio of 4:1, followed by reconstitution for at least 2 months. Mice were bred and maintained in a specific pathogen-free facility in the Department of Comparative Medicine of the Mayo Clinic. All animal protocols were approved by the Institutional Animal Care and Use Committees (IACUC) of the Mayo Clinic Rochester.

### Cell lines

The retroviral packaging Plat-E cells were a gift from Dr. Hongbo Chi (St. Jude Children’s Research Hospital) and from female origin. The cells were cultured in Dulbecco’s Modified Eagle Medium (DMEM, Thermo Fisher Scientific) supplemented with 10% fetal bovine serum (FBS) and 2 mM glutamine, 100 U/ml Penicillin and 100 mg/ml Streptomycin and maintained at 37°C in 5% CO2. Puromycin (1 mg/ml) and blasticidin (10 mg/ml) antibiotics were also added into the culture medium to maintain selective pressure and were removed one day before retrovirus plasmid transfection.

### Immunizations and other mouse experimentation

For tamoxifen treatment, mice were injected intraperitoneally with tamoxifen (1 mg per mouse) in corn oil daily for four consecutive days and analyzed 8 days after the last injection. For experiments involving an immune challenge, chimeras were given tamoxifen (1 mg per mouse) in corn oil daily for four consecutive days via oral gavage and challenged with antigens or influenza at day 8 after the last tamoxifen administration. For NP-OVA immunization experiments, antigen for immunization was prepared by mixing NP_20_-OVA (20 molecules of NP linked to OVA; Biosearch Technologies), 10% KAL(SO_4_)_2_ dissolved in PBS at a ratio of 1:1, in the presence of LPS (Escherichia coli strain 055:B5; Sigma) at pH 7. Mice were immunized intraperitoneally (100 μg NP-OVA and 10 μg LPS) for analysis of NP-specific antibody response. Nine days after immunization, sera, spleens and mesenteric lymph nodes were collected from the mice. Thirteen days after infection, spleens, mediastinal lymph nodes and lungs from the mice were harvested for analysis. For T-independent immune response, mice were given 50 μg TNP-LPS (Biosearch Technologies) intraperitoneally. Sera and spleens were collected on day 7 after immunization.

### Cell isolation and culture

Mouse B cells were isolated from pooled single cell suspensions of spleen and peripheral lymph nodes using CD19 microbeads (Miltenyi, catalog no. 130-052-201) or EasySep Mouse B Cell Isolation Kit (Stemcell Technologies, catalog no. 19854). B cells were cultured in RPMI1640 medium supplemented with 10% (vol/vol) FBS and 1% penicillin-streptomycin and activated with 3 μg/mL LPS (Sigma-Aldrich), 10 ng/mL recombinant mouse IL-4 (Tonbo Bioscience) plus 20 ng/mL recombinant human BAFF (Biolegend) for 3 days. Alternatively, B cells were activated with10 μg/mL anti-IgM (Jackson Immunoresearch), 5 μg/mL anti-CD40 (Bio X Cell) and 10 ng/mL recombinant mouse IL-4, or 2.5 μM CpG ODN2006 (Integrated DNA Technologies) together with 10 ng/mL recombinant mouse IL-4 or 3 μg/mL LPS with 10 ng/mL recombinant murine IFN-γ (Peprotech) and 20 ng/mL recombinant human BAFF or 5 μg/mL anti-CD40 (Bio X Cell) and 10 ng/mL recombinant mouse IL-4. B cell proliferation was measured by CellTrace violet dye dilution (Thermo Fisher Scientific).

### Retroviral constructs and transductions

WT TFEB retroviral constructs were made by inserting the cDNA of mouse TFEB isoform b into the MSCV-IRES-EGFP retroviral vector. Ca TFEB retroviral constructs were generated by mutating mouse TFEB isoform b at Ser142 and Ser211 to Ala, and the mutated form was inserted into the MSCV-IRES-EGFP vector. TFEB TDN retroviral construct was generated by deleting activation domain (AD), binding region (BR), and proline-rich region, while retaining bHLH and Leucine-zipper region. HA-tag and nuclear location sequence were added at C terminal of the construct. Mito-QC reporter retroviral construct was a gift from Dr. Ping-Chih Ho (Ludwig Institute for Cancer Research), as described previously(*56*). To obtain infectious retroviral stocks, each construct was transfected into Plat-E cells along with pCL-Eco packaging plasmid using Lipofectamine™ 3000 Transfection Reagent. Spleen B cells were activated with 3 μg/mL LPS, 10 ng/mL recombinant mouse IL-4, or 0.25 μg/ml anti-CD180 for 40 h, followed by transduction with indicated viruses in the presence of 8 μg/ml polybrene at 32°C, 2000 rpm for 90 min. Transduced B cells were put in the 37°C incubator for 4 hours, then changed to fresh medium with 3 μg/mL LPS, 10 ng/mL recombinant mouse IL-4, and 20 ng/mL recombinant human BAFF for further activation. Transduced cells were analyzed after 3 days of activation.

### Flow cytometry

For analysis of surface markers, cells were stained in phosphate-buffered saline (PBS) containing 1% (w/v) bovine serum albumin (BSA) with indicated antibodies. The following antibodies were used: anti-B220 (RA3-6B2), anti-CD19 (6D5), anti-TCRβ (H57-597), anti-CD24 (M1/69), anti-BP-1 (6C3), anti-IgD (11-26c.2a), anti-IgG1 (RMG1-1), anti-CD25 (PC61), anti-CD4 (RM4-5), anti-CD21 (7E9), anti-CD23 (B3B4), anti-CD93 (AA4.1), anti-CD138 (281–2), anti-GL7 (GL7), anti-CD278 (C398.4A), anti-PD-1 (J43), anti-CD86 (GL-1), anti-IL-7Ra (A7R34), and anti-CD71 (R17217) were all from Biolegend, anti-IgM (II/41) and anti-CD184 (2B11) were purchased Thermo Fisher Scientific. Anti-CD95 (Jo2), and anti-CD43 (S7) were obtained from BD Biosciences. CXCR5 was stained with biotinylated anti-CXCR5 (2G8) and streptavidin-conjugated PE (both from BD Biosciences) to enhance the signal. Intracellular Foxp3 (FJK-16s), Ki-67 (SolA15, Thermo Fisher Scientific), and anti-Bcl6 (K112-91, BD Biosciences) were analyzed in cells fixed and permeabilized with Foxp3 staining buffers according to the manufacturer’s instructions (Thermo Fisher Scientific). For phosphoflow staining, cells were stained with surface markers, then fixed with 1× Lyse/Fix (BD Biosciences) buffer at 37°C for 10 min, washed and permeabilized by ice-cold Perm III buffer (BD Biosciences) on ice for 30 min, followed by staining with anti-phospho-S6 (S235/236) or anti-phospho-4E-BP1 (T37/46) (both from Cell Signaling Technology) for 30 min at room temperature. Cell viability was examined by Fixable viability dye (Tonbo Bioscience) or 7-AAD (Thermo Fisher Scientific) following the manufacturer’s protocol. For LAMP1 staining, surface staining was done with FACS buffer on ice, followed by fixation and permeabilization using BD Cytofix/Cytoperm Fixation/Permeabilization Kit (BD Biosciences), LAMP1 antibody (1D4B, BioLegend) was diluted in BD Perm/Was Buffer and stained at room temperature for 30 min. For the dye staining, B cells were stained with 20 nM MitoTracker Deep Red (ThermoFisher Scientific), 20 nM MitoTracker Green (ThermoFisher Scientific), 100 nM tetramethylrhodamine methyl ester (TMRM, ThermoFisher Scientific). 500 nM CellROX or 1 μM MitoSOX in HBSS at 37°C for 20 min. Flow cytometry was performed on a BD Fortessa X-20 or LSR II instrument or Attune NxT system (Life Technologies). Data were then analyzed by FlowJo software (Tree Star).

### p-S65-Ub sandwich ELISA

Levels of phosphorylated ubiquitin at serine 65 (p-S65-Ub) were assessed in a sandwich type ELISA on a Meso Scale Discovery (MSD) platform that uses electrochemiluminescence (ECL) as a readout, which was slightly modified from *Watzlawik et al., 2021*(*67*). In brief, here we used a SULFO-TAG-labeled mouse anti-Ub antibody (clone P4D1) instead of two subsequent detecting antibodies (1. Ub (P4D1) followed by 2. SULFO-TAG-labeled anti-mouse antibody) as described previously(*67*).

#### A. SULFO-TAG-labeling

For SULFO-TAG-labeling of the Ub detecting antibody (ThermoFisher #14-6078-37, Ub (clone P4D1)), we first removed sodium azide by using Amicon ultra 0.5 ml centrifugal filters with a 50 kDa MWCO (Millipore, UFC505008) and washed 5 times with PBS, pH: 7.9 at 14,000 × g for 2 minutes. Sample recovery was done by inverting the filter in a new tube and spinning for another 2 minutes at 1000 × g. Ub (P4D1) antibody was then incubated at room temperature for 2 h with SULFO-TAG NHS-Ester (MSD, #R31AA) in a challenge ratio of 20:1 (Sulfotag NHS-Ester: antibody) on a rotational shaker. Excess, non-conjugated SULFO-TAG was removed by using a 0.5 ml Zeba Spin desalting column (40K MWCO) (ThermoFisher, A57760) according to the manufacturer’s recommendation.

#### B. p-S65-Ub ELISA

p-S65-Ub antibody (CST #62802) was used as a capture antibody in a concentration of 1 μg/ml in 200 mM sodium carbonate buffer pH 9.7 and coated overnight at 4°C with 30 μl per well in 96-well MSD plate (MULTI-ARRAY® 96-well Plate; L15XA-3). The next morning MSD plates were washed 2 times with 0.22-micron filtered ELISA wash buffer (150 mM Tris, pH 7.4, 150 mM NaCl, 0.1% [v:v] Tween-20) and subsequently blocked by adding ELISA blocking buffer (150 mM Tris, pH 7.4, 150 mM NaCl, 0.1% [v:v] Tween-20, 1% BSA [w:v]) and incubated for 1 h at 22°C without shaking. All samples were run in duplicates and diluted in blocking buffer using 10 μg of total protein per well. Antigens were incubated for 2 h at 22°C on a microplate mixer (USA Scientific, 8182-2019) at 500 rpm and three washing steps were then performed as described before. SULFO-TAG-labeled Ub (P4D1) antibody (1 µg/ml) was added in blocking buffer in 50 μl total volume per well and incubated for 2 h at 22°C on a microplate mixer at 500 rpm. After three washing steps, 150 μl MSD GOLD Read Buffer (R92TG-2) were finally added to each well and the plate being read on a MESO QuickPlex SQ 120 reader.

### Metabolic assays

The bioenergetic activities of B cells, displayed by both ECAR and OCR were measured by Seahorse assays according to the established protocols from Agilent Technologies. Briefly, B cells were seeded at 150, 000-300, 000 cells/well on Cell-Tak (Corning) coated XFe96 plate in indicated medium (For OCR: Seahorse XF RPMI medium containing 10 mM glucose, 2 mM L-glutamine, and 1 mM sodium pyruvate, pH 7.4; For ECAR: Seahorse XF RPMI medium plus 2 mM L-glutamine, pH 7.4; all reagents from Agilent Technologies). For the Mitostress test, OCR and ECAR were measured in the presence of Oligomycin (1.5 μM, Sigma-Aldrich), FCCP (1.5 μM, Sigma-Aldrich), and Rotenone (1 μM, Sigma-Aldrich)/ Antimycin A (1 μM, Sigma-Aldrich). For glycolysis stress, both OCR and ECAR were measured by sequential injection of Glucose (10 mM, Agilent Technologies), Oligomycin (1.5 μM, Sigma-Aldrich), 2-DG (50 mM, Sigma-Aldrich). Glycolytic flux was also measured by detritiation of [3-^3^H]-glucose (Perkin Elmer) as described(*72*). Briefly, 1 μCi [3-^3^H] glucose was added into the culture media, and 2 h later, 500 μL media were transferred to a 1.5 mL microcentrifuge tube containing 50 μL of 5 N HCl. The microcentrifuge tubes were then placed in 20 mL scintillation vials containing 0.5 ml water with the vials capped and sealed. ^3^H_2_O was separated from unmetabolized [^3^H] glucose by evaporative diffusion for 24 h at room temperature. A cell-free sample containing 1 μCi ^3^H-glucose was included as a background control.

### Transmission electron microscopy

B cells were cultured with LPS, IL-4 plus BAFF for 3 days, and cell pellets were harvest and fixed in 2% paraformaldehyde and 2.5% glutaraldehyde in 0.1 M sodium cacodylate. Following fixation, cells were embedded and sliced for transmission electron microscopy. The Grids were imaged with a JEM1400 plus transmission electron microscope (JEOL). Damaged mitochondria with mitophagy were defined as either no visible cristae, surrounded by a phagophore or being located inside an amphisome.

### Single cell RNA sequencing analysis

The primary data was obtained from ArrayExpress (E-MTAB-9478), followed by annotation and alignment using CellRanger. All samples from spleen were included. Events with 200-5000 genes detected per cell (nFeature) and <5% mitochondrial genes were put through further analysis with the package of Seurat (v4) in R project. Our methodology for analysis was adapted from the original vignette (https://satijalab.org/seurat/articles/integration_introduction.html). In brief, differential expressed genes were found in each dataset using Principal Component Analysis. Shared genes were then identified across different time points as “anchors” to integrate all datasets. Subsequently, clustering was performed based on K-nearest neighbor (KNN) graph constructed. Clusters resembled contaminating cells (i.e., T cells, myeloid cells etc.) were excluded. The data was re-clustered using the workflow described previously.

For cell function evaluation, the “AddModuleScore ()” function was employed. Gene sets displayed in Fig.1h-i included the following: Hallmark mTORC1 signaling (MSigDB, M5924), Hallmark oxidative phosphorylation (MSigDB, M5936), Putative TFEB target genes (described previously(*61*)), KEGG Lysosome (MSigDB, M11266), GO lysosome localization (GO:0032418), mitochrondrial complex I (MSigDB, M39781), Reactome mitophagy (MSigDB, M27418), and Reactome PINK1-PRKN mediated mitophagy (MSigDB, M27419).

### Bulk RNA sequencing

WT or RagA/RagB deficient B cells were activated *in vitro* for 72 h with LPS, IL-4, and BAFF. RNA was isolated using a Quick-RNA Microprep kit (ZYMO research) following the manufacturer’s instructions. After quality control, high quality total RNA was used to generate the RNA sequencing library. Reads with low quality, containing the adaptor (adaptor pollution), or with high levels of N base were removed to generate clean data. HISAT(*73*) was used to align the clean reads to the mouse reference genome (Mus_musculus, GCF_000001635.27_GRCm39). Bowtie2(*74*) was used to align the clean reads to the reference genes. DEG analysis was carried out using DESeq2, and genes with log2FC > 0 and false discovery rate < 0.05 were considered for gene cluster analysis.

### Nuclear and Cytoplasmic Extraction

Two million B cells were collected and washed with ice-cold PBS twice, discarded the supernatant, and collected the cell pellets. Gently resuspend cells in 200 μl 1× Hypotonic Buffer (20 mM Tris-HCl, pH 7.4, 10 mM NaCl, 3 mM MgCl_2_) by pipetting up and down several times, then incubated on ice for 15 minutes. 15 μl detergent (10% NP40) was added into the suspension and vortexed for 10 seconds at the highest speed. The supernatant was collected after centrifuging for 10 min at 5000 rpm at 4°C. This supernatant contains the cytoplasmic fraction. The collected cell pellets were washed once with 1× Hypotonic Buffer and resuspended in 50 μl Complete Cell Extraction Buffer (10 mM Tris, pH 7.4, 2 mM Na_3_VO_4_, 100 mM NaCl, 1% Triton X-100, 1 mM EDTA, 0% glycerol, 1 mM EGTA, 0.1% SDS, 1 mM NaF, 0.5% deoxycholate, 20 mM Na_4_P_2_O_7_) for 30 min on ice with vortexing at 10 min intervals. The supernatant (nuclear fraction) was collected after centrifuging for 30 min at 14,000 g at 4°C, and the nuclear extracts were ready for assay.

### Immunoblots

B cells were lysed in radioimmunoprecipitation assay (RIPA) buffer (50 mM Tris (pH 7.4), 150 mM NaCl, 1% NP-40, 0.5% sodium deoxycholate, 0.1% sodium dodecyl sulfate (SDS)) supplemented with protease inhibitor cocktail and phosphatase inhibitor cocktail (Sigma-Aldrich). Protein concentration was detected by BCA assay (Thermo Fisher Scientific), and an equal amount of protein was resolved in 4-12% SDS-polyacrylamide gel electrophoresis (SDS-PAGE) (Bio-Rad). Proteins were transferred to polyvinylidene difluoride membranes (Millipore) and probed overnight with the following primary antibodies: anti-p-S6K (108D2), anti-p-S6 (D57.2.2E), anti-LAMP1(C54H11), anti-p-4EBP1 (236B4), anti-Raptor (24C12), and anti-RagA (D8B5) all from Cell Signaling Technology), anti-AID (mAID-2, Thermo Fisher Scientific), anti-TFEB (A303-673A, Bethyl Laboratories), Lamin B (66095-1-Ig, Proteintech), tubulin (11224-1-AP, Proteintech), TFE3 (HPA023881, Sigma-Aldrich) and anti-β-actin (13E5, Sigma-Aldrich). The membrane was washed and incubated with indicated secondary antibody for the subsequently enhanced chemiluminescence (ECL, Thermo Fisher) exposure.

### Immunofluorescence

For observing the germinal center structure in the spleen, part of the spleen from the immunized or infected mice was fixed in 4% Paraformaldehyde at 4°C overnight, then the spleens were dehydrated in 20% sucrose for at least 36 h. Then the spleens were embedded in OCT and sectioned at 5 μm thickness. The slides were air-dried at room temperature before being fixed in cold acetone at −20 °C for 10 min. The fixed slides were washed with PBS for twice and blocked with Blocking buffer (Thermo Fisher Scientific) for 1 h at room temperature. Biotinylated Peanut Agglutinin (PNA, Vector laboratories) was stained o44n the sections at 4°C overnight. The slides were washed with PBST 3 times, then the Alexa Fluor™ 488 Conjugated Streptavidin, anti-mouse CD21/CD35 (Alexa Fluor® 594, Biolegend), and anti-mouse IgD (Alexa Fluor® 647 anti-mouse IgD, Biolegend) was applied onto the section for 2 h at room temperature. After washing with PBST for 3 times, the slides were counterstained with 4’, 6-diaminodino-2-phenylindole (DAPI) and mounted. The stained slides were reviewed, and representative images were acquired on Olympus DP80 digital microscope.

### RNA Isolation and Real-time Quantitative PCR

Total RNA was extracted using the RNeasy Micro kit (Qiagen) according to the manufacturer’s instructions, and total RNA was reverse transcribed into cDNA by PrimeScript RT Reagent Kit (Takara) following the established protocol of the kit. The mRNA level of *Rragd, Rragc, Bhlhe40,* and *Prodh2* was detected by real-time PCR with a Thermo Fisher Real-time PCR system, while β-actin was used as an internal control. Each sample was analyzed in triplicate and the relative amount of gene expression was calculated using the 2^−ΔΔCt^ method.

### ELISA

For detecting NP-specific antibodies in sera, wells were coated with 1 μg/mL NP_23_-BSA or NP_2_-BSA in coating buffer (Bicarbonate-carbonate buffer, pH 9.6) overnight. Plates were washed twice with washing buffer (0.05% Tween 20 in PBS), blocked with 5% blocking protein (Bio-Rad) at 37°C for 1 h, washed twice, and incubated with indicated sera samples at 37°C for 1.5 h.

Horseradish peroxidase (HRP)-conjugated secondary antibodies: anti-mouse IgG1, anti-mouse IgG2b, anti-mouse IgG2c, and anti-mouse IgG3 (all from SouthernBiotech), or anti-mouse IgM (Bethyl laboratories) were developed at 37°C for 1 h after washing with a buffer for four times. The reaction was further developed with tetramethylbenzidine (TMB), then stopped by 2N H_2_SO_4_, and read at 450 nm. Sera from influenza-challenged mice were assayed using total viral lysate prepared from stocks of strain Influenza A/PR8/34.

### Preparation of amino acid medium

Amino acid-free (AA-) medium was prepared by RPMI 1640 powder (R8999-04A, US Biological Life Science) and sodium phosphate dibasic (5.6 mM, the same concentration as commercially available RPMI 1640 medium, US Biological Life Science), supplemented with 10% (v/v) dialyzed FBS (Thermo Fisher Scientific). Amino acid-sufficient (AA+) medium was prepared by adding proper volumes of MEM amino acids solution (essential amino acids, EAA, 50×), MEM non-essential amino acids solution (NEAA, 100×), and 200 mM L-Gln (all from Sigma-Aldrich) to AA-medium to reach a final concentration of 1×EAA, 1×NEAA, and 2 mM Gln. The medium was supplemented with 10% (v/v) dialyzed FBS. Medium containing single amino acids (Ala, Leu, Gln, or Arg) or their combinations was prepared with AA– medium (prepared to the same concentrations present in the AA+ medium). All media were adjusted to pH7.5 and filter-sterilized (0.2 μm) before use.

### Statistical analysis

Statistics were performed on GraphPad Prism 8. P values were calculated with Student’s t-test, one-way ANOVA, or two-way ANOVA, as indicated in the figure legends. p < 0.05 was considered significant. All error bars were represented as SEM.

## Acknowledgements

We thank Drs. Andrea Ballabio, Ming O. Li, Ping-Chih Ho for sharing mouse strains and research reagents. We acknowledge the Microscopy and Cell Analysis Core at Mayo Clinic Rochester. We acknowledge the NIH (grants R01 AI 162678 and R01 AR077518) for supporting this work in H.Z.’s laboratory, and RO1 AI154598, RO1 AI147394, RO1 AI176171 and RO1 AI 112844 to J.S.’s laboratory.

## Author contributions

X.X.Z. and H.Z. conceived the project, designed the research, interpreted the data and wrote the manuscript. X.X.Z. and X.Z. prepared the materials and carried out the experiments. Y.W. and Y.M.C. performed the bioinformatics analysis. Y.L. managed the mouse colony, performed molecular biology experiments and fluorescence imaging. D.B. provided imaging analysis. V.S. provided antibodies and other research materials. J.O.W. and W.S. performed part of the mitophagy analyses. A.L.R. provided tissue samples from Raptor mutant mouse line. M.R.B. interpreted the data and revised the manuscript. This work is a collaboration with J.S. and M.R.B., who provided key materials and expertise for the research.

---

**Figure S1.**
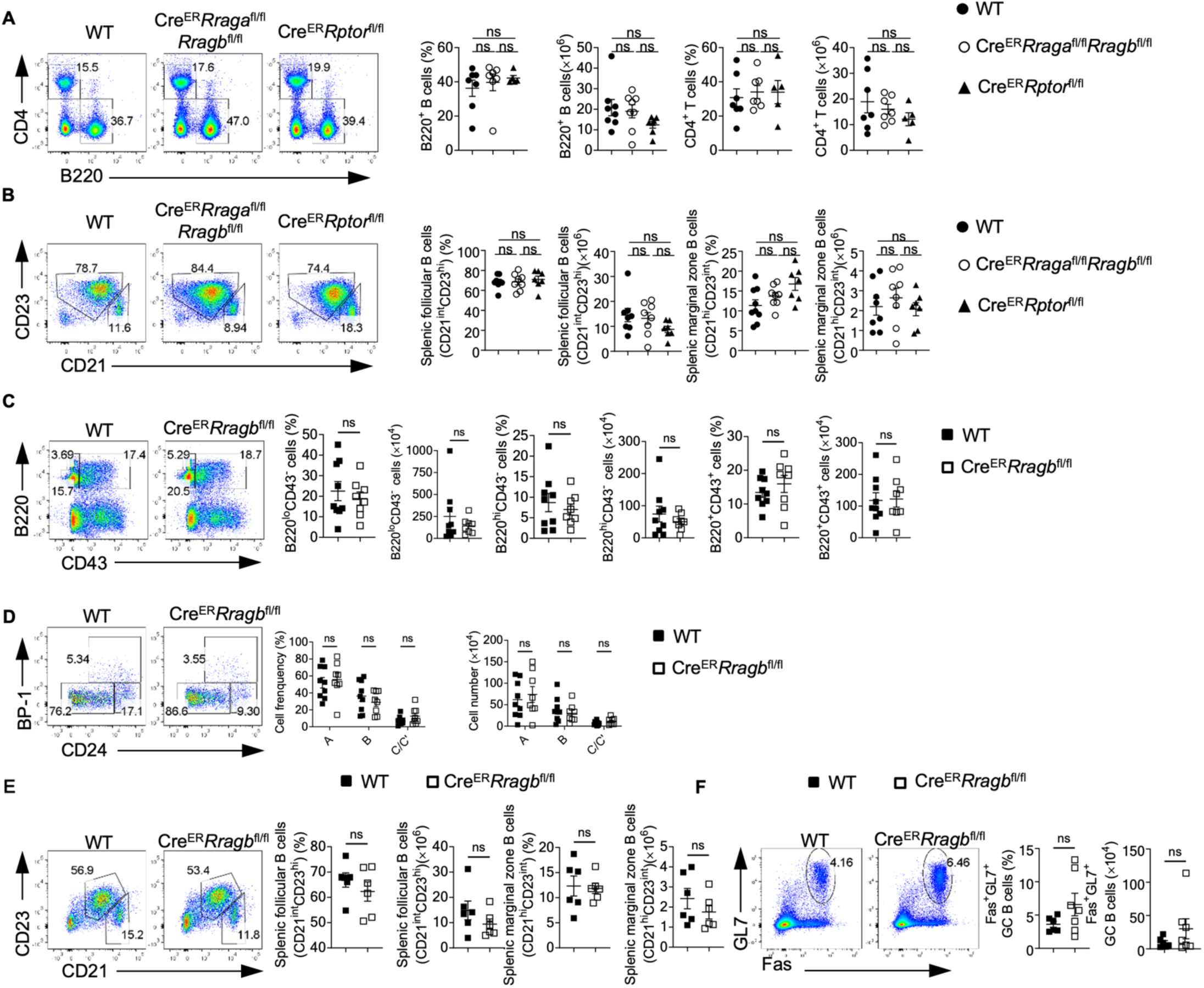
RagB deficiency doesn’t affect early B cell development in BM and peripheral B cells. (A) Representative flow plots of splenic CD4^+^ T cells and B220^+^ B cells from WT mice (n = 7), *Cre^ER^Rraga*^fl/fl^*Rragb*^fl/fl^ (n = 7), and *Cre^ER^Rptor*^fl/fl^ (n = 5). Right, summaries of the percentages and numbers of splenic CD4^+^ T cells (gated on TCR-b^+^ cells) and B220^+^ B cells. (B) flow plots of splenic follicular B cells (CD21^int^CD23^hi^) or marginal zone (CD21^hi^CD23^int^) from WT mice (n = 8), *Cre^ER^Rraga*^fl/fl^*Rragb*^fl/fl^ (n = 8), and *Cre^ER^Rptor*^fl/fl^ (n = 7). Right, summaries of the percentages and numbers of splenic follicular B cells (CD21^int^CD23^hi^) or marginal zone (CD21^hi^CD23^int^). (C) Representative flow plots of bone marrow B220 and CD43 expression from WT (n = 9), and Cre^ER^*Rragb*^fl/fl^ (n = 8) mice. Right, summaries of the percentages and numbers of B220^lo^CD43^−^, B220^hi^CD43^−^ and B220^+^CD43^+^ cells. (D) Representative flow plots of BP-1 and CD24 expression in BM B220^+^CD43^+^IgM^−^ B cell precursors from WT (n = 9) and Cre^ER^*Rragb*^fl/fl^ (n = 8) mice. Right, summary of the percentages and numbers of fraction A (CD24^−^BP-1^−^), fraction B (CD24^+^BP-1^−^), and fraction C/C′(CD24^+^BP-1^+^) cells. (E) Representative flow plots of splenic follicular B cells (CD21^int^CD23^hi^) or marginal zone (CD21^hi^CD23^int^) from WT (n = 9) and Cre^ER^*Rragb*^fl/fl^ (n = 8) mice. Right, summaries of the percentages and cell numbers of splenic follicular B cells (CD21^int^CD23^hi^) or marginal zone (CD21^hi^CD23^int^) B cells. (F) Representative flow plots of GL-7 and Fas expression in lymphocytes from the Peyer’s patches from WT (n = 6) and Cre^ER^*Rragb*^fl/fl^ (n = 7) mice. Right, summary of the percentages and numbers of GC (GL-7^+^Fas^+^) B cells. Data in graphs represent mean ± SEM. ns, not significant. One-way ANOVA (A and B), two-way ANOVA (D), two-tailed Student’s t test (C, E and F).

**Figure S2.**
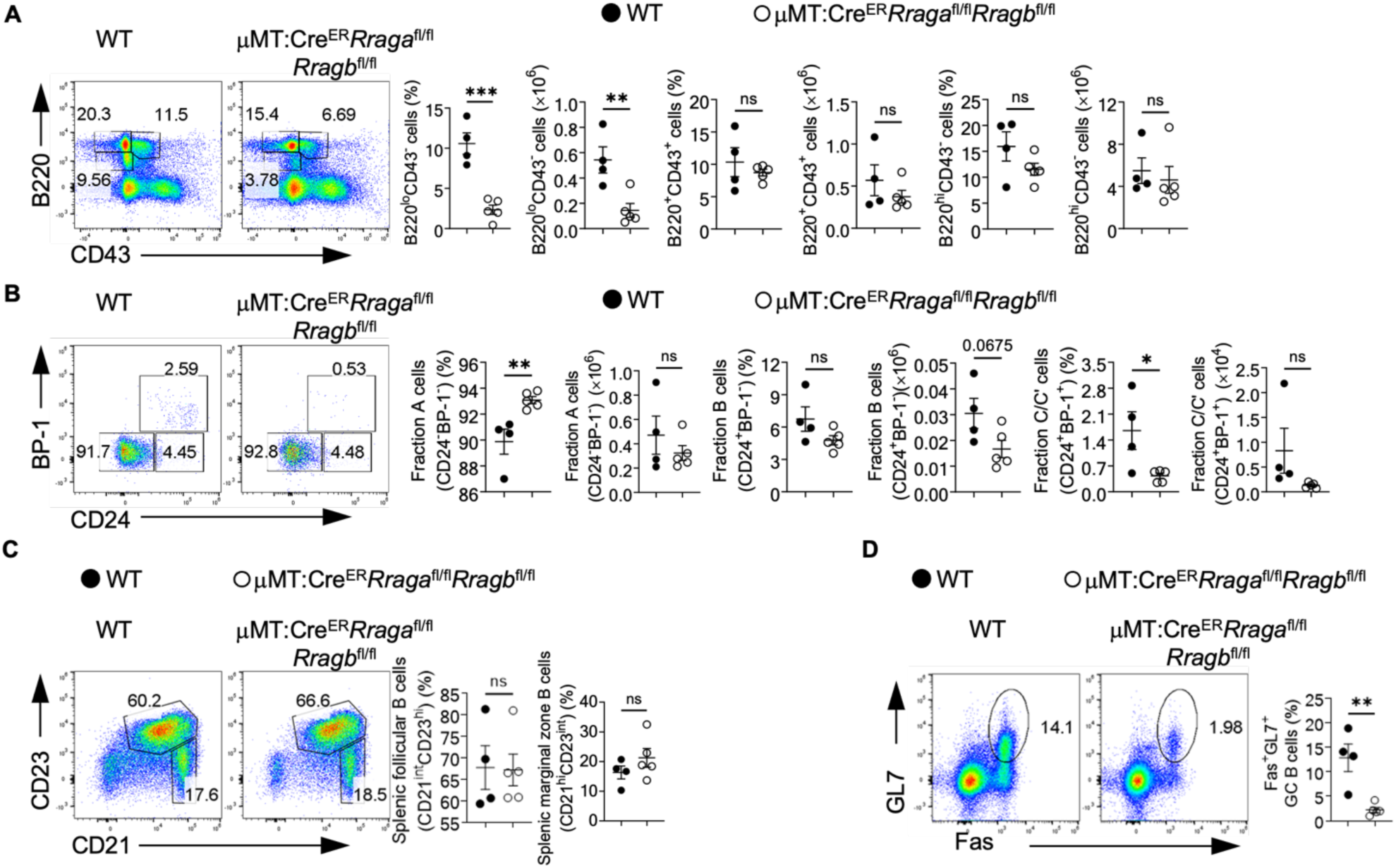
Rag-GTPases deficiency intrinsically disrupts B cell development. (A-D) Tamoxifen was administered to μMT:Cre^ER^*Rraga*^fl/fl^*Rragb*^fl/fl^ (n = 5), and WT (n = 4) chimera mice by oral gavage daily for 4 consecutive days. Mice were analyzed 7 days after the last injection. (A) Representative flow plots of bone marrow B220 and CD43 expression from μMT:Cre^ER^*Rraga*^fl/fl^*Rragb*^fl/fl^, and WT chimera mice. Right, summaries of the percentages and numbers of B220^lo^CD43^−^, B220^hi^CD43^−^ and B220^+^CD43^+^ cells. (B) Representative flow plots of BP-1 and CD24 expression in BM B220^+^CD43^+^IgM^−^ B cell precursors from μMT:Cre^ER^*Rraga*^fl/fl^*Rragb*^fl/fl^, and WT chimera mice. Right, summaries of the percentages and numbers of bone marrow fraction A (CD24^−^BP-1^−^), fraction B (CD24^+^BP-1^−^), and fraction C/C′(CD24^+^BP-1^+^) cells. (C) Representative flow plots of splenic follicular B cells (CD21^int^CD23^hi^) or marginal zone (CD21^hi^CD23^int^) from μMT:Cre^ER^*Rraga*^fl/fl^*Rragb*^fl/fl^, and WT chimera mice. Right, summaries of the percentages of splenic follicular B cells (CD21^int^CD23^hi^) or marginal zone (CD21^hi^CD23^int^) B cells. (D) Representing flow plots of Fas and GL-7 expression on B cells from the Peyer’s patches of μMT:Cre^ER^*Rraga*^fl/fl^*Rragb*^fl/fl^, and WT chimera mice. Right, summary of the percentages of GC (GL-7^+^Fas^+^) B cells. Data in graphs represent mean ± SEM. ns, not significant. *p < 0.05, **p < 0.01, ***p < 0.001, two-tailed Student’s t test was used.

**Figure S3.**
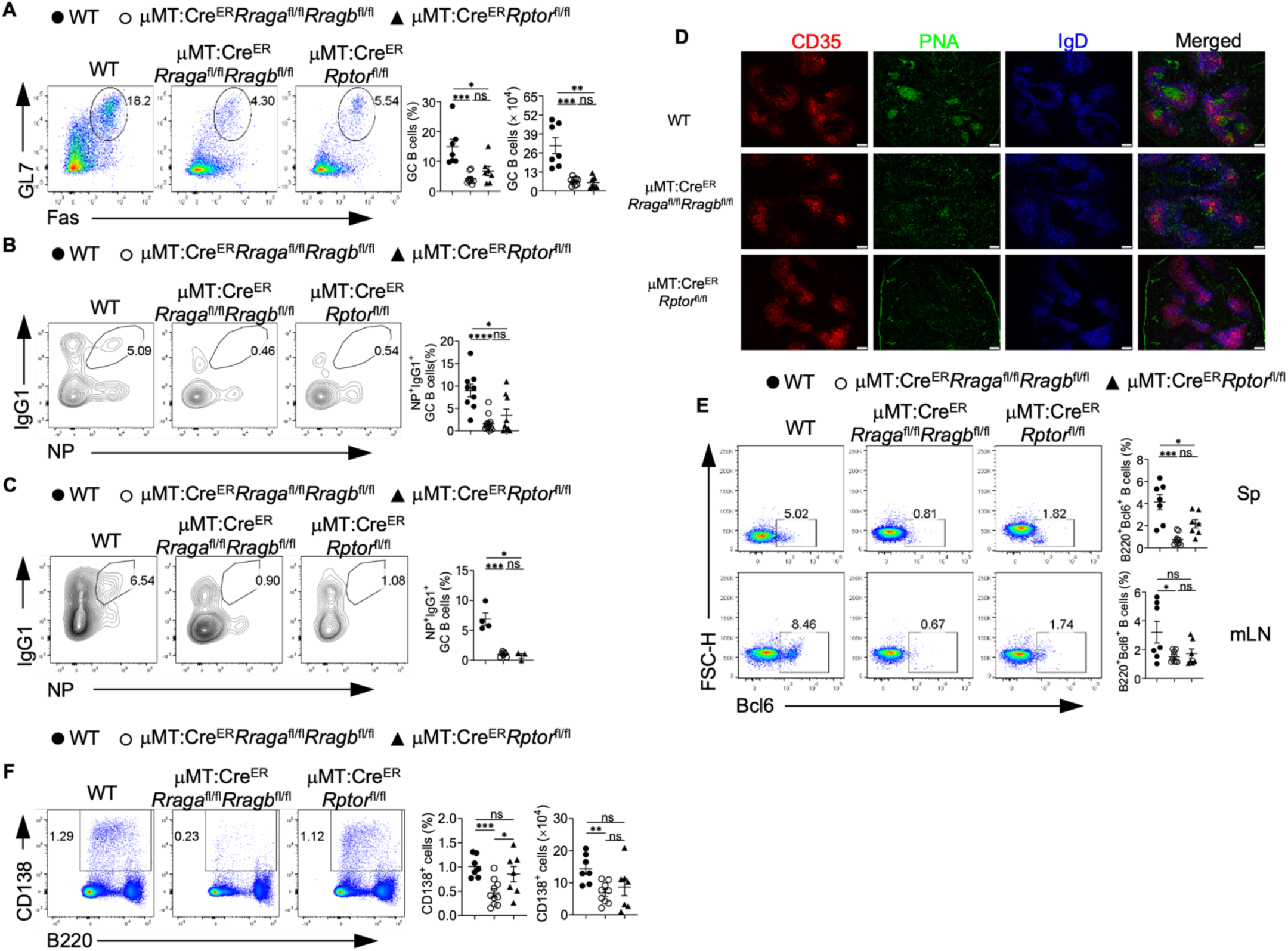
Rag-GTPases deficiency reduces GC formation and antibody production independent of mTORC1 in peptide immunization. (A-F) Tamoxifen was administered to μMT:Cre^ER^*Rraga*^fl/fl^*Rragb*^fl/fl^, μMT:Cre^ER^*Rptor*^fl/fl^, and WT chimera mice by oral gavage daily for 4 consecutive days. Mice were immunized intraperitoneally (100 mg NP-OVA/alum) 7 days after the last injection. (A) Representative flow plots of GL-7 and Fas expression on B cells from the mesenteric lymph node (mLN) of immunized WT (n = 7), μMT:Cre^ER^*Rraga*^fl/fl^*Rragb*^fl/fl^ (n = 9), and μMT:Cre^ER^*Rptor*^fl/fl^ (n = 7) chimera mice. Right, summary of the percentages and numbers of GC (GL-7^+^Fas^+^) B cells. (B) Representative flow plots of NP^+^IgG1^+^ GC B cells from the spleen of immunized WT (n = 9), μMT:Cre^ER^*Rraga*^fl/fl^*Rragb*^fl/fl^ (n = 13), and μMT:Cre^ER^*Rptor*^fl/fl^ (n = 9) chimera mice. Right, summary of the percentages of NP^+^IgG1^+^ GC B cells. (C) Representative flow plots of NP^+^IgG1^+^ GC B cells from the mLN of immunized WT (n = 4), μMT:Cre^ER^*Rraga*^fl/fl^*Rragb*^fl/fl^ (n = 7), and μMT:Cre^ER^*Rptor*^fl/fl^ (n = 3) chimera mice. Right, summary of the percentages of NP^+^IgG1^+^ GC B cells. (D) Cryosections from the spleens of immunized WT, μMT:Cre^ER^*Rraga*^fl/fl^*Rragb*^fl/fl^, and μMT:Cre^ER^*Rptor*^fl/fl^ chimera mice were stained for GC B cells (anti-PNA). Scale bar, 50 mm. Anti-IgD stains for FoBs, anti-CD35 stains for follicular dendritic cells (LZ). (E) Representative flow plots of Bcl6 expression on total B cells from both spleen (upper panel) and mLN (lower panel) of immunized WT (n = 7), μMT:Cre^ER^*Rraga*^fl/fl^*Rragb*^fl/fl^ (n = 9), and μMT:Cre^ER^*Rptor*^fl/fl^ (n = 7) chimera mice. Right, summary of the percentage of Bcl6^+^ B cells on B cells. (F) Representative flow plots of CD138 and B220 expression in mLN of WT (n = 7), μMT:Cre^ER^*Rraga*^fl/fl^*Rragb*^fl/fl^ (n = 9), and μMT:Cre^ER^*Rptor*^fl/fl^ (n = 7) mice. Right, summary of the percentages and numbers of CD138^+^ cells. Data in graphs represent mean ± SEM. ns, not significant. *p < 0.05, **p < 0.01, ***p < 0.001, and ****p < 0.0001, one-way ANOVA was used.

**Figure S4.**
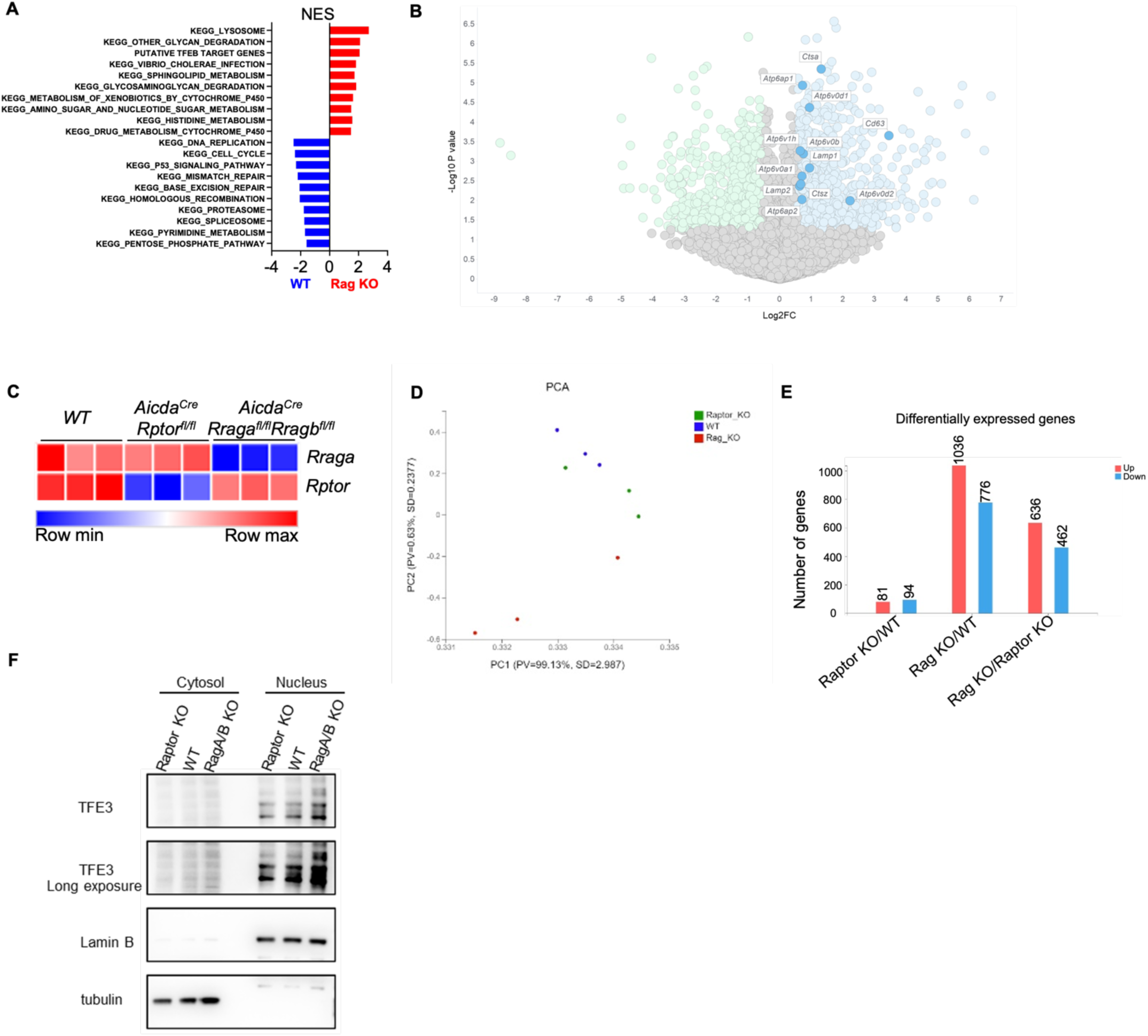
Rag-GTPases regulate TFEB/TFE3 axis in B cells. (A) Bulk RNA sequencing was performed on the B cells activated with LPS/IL-4/BAFF for 72 h, then GSEA was conducted on differentially expressed genes (DEGs), and significantly enriched pathways were plotted according to NES (normalized enrichment score). (B) Volcano plot of the lysosome genes enriched in RagA/RagB deficient B cells. (C-E) GC B cells were sorted from immunized *Aicda^Cre^Rraga*^fl/fl^*Rragb*^fl/fl^, *Aicda^Cre^Rptor*^fl/fl^, and WT mice, and conducted bulk RNA sequencing. (C) *Rraga,* and *Rptor* levels from the bulk RNA sequencing data. (D) PCA plot from the bulk RNA sequencing analysis. (E) Statistics of differently expressed genes from different comparisons. (F) Cytosolic and nuclear proteins were isolated from the B cells activated with LPS/IL-4/BAFF for 72 h. Expression of TFE3 was examined by immunoblot. Lamin B was used as a nuclear control, tubulin was used as a cytosol control.

**Figure S5.**
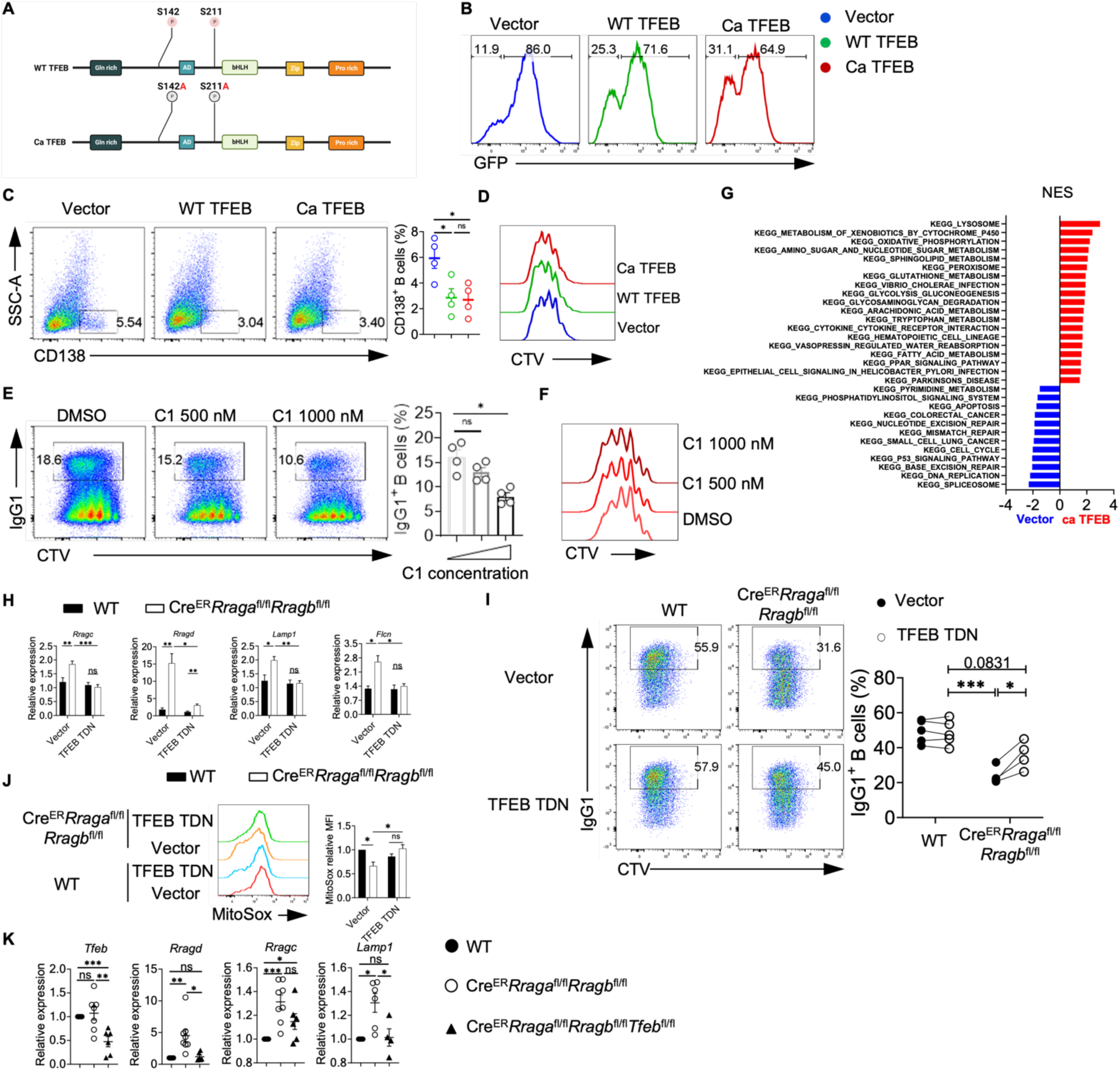
TFEB plays a critical role in Rag GTPases regulated B cell function. (A) Schematic of the experimental design for making WT TFEB and Ca TFEB plasmids. (B) Representative histogram of GFP expression on the vector, WT TFEB or Ca TFEB transduced B cells. (C) Representative flow plots of CD138^+^ expression on vector, WT TFEB, and Ca TFEB transduced GFP^+^ B cells. (D) Representative flow plots of CTV dilution from vector, WT TFEB, and Ca TFEB transduced GFP^+^ B cells. (E) Representative flow plots of IgG1 and CTV expression activated in LPS/IL-4/BAFF with different concentrations of compound C1 for 72 h. Right, summary of IgG1^+^ B cells. (F) Representative flow plots of CTV dilution from different concentrations of compound C1 treated B cells. (G) GFP^+^ B cells were sorted from vector, or Ca TFEB transduced B cells, and bulk RNA sequencing was conducted on these samples. GSEA was performed on the DEGs, and significantly enriched pathways were plotted according to NES. (H) Splenic B cells from Cre^ER^*Rraga*^fl/fl^*Rragb*^fl/fl^ (n = 5), and WT (n = 5) mice were transduced with vector or TFEB TDN, then B cells were cultured with LPS/IL-4/BAFF for 72 h, TFEB downstream target genes were examined. (I) Spenic B cells from Cre^ER^*Rraga*^fl/fl^*Rragb*^fl/fl^ (n = 4), and WT (n = 5) mice were transduced with vector or TFEB TDN, and IgG1^+^ B cells were measured. Right, summary of the IgG1^+^ B cells. (J) Spenic B cells from Cre^ER^*Rraga*^fl/fl^*Rragb*^fl/fl^ and WT mice were transduced with vector or TFEB TDN, MitoSox was measured after activation with LPS/IL-4/BAFF for 72 h. Right, summary of MitoSox level (Relative to vector transduced WT B cells). (K) TFEB downstream target genes were measured on the activated B cells from WT (n = 4-6), Cre^ER^*Rraga*^fl/fl^*Rragb*^fl/fl^ (n = 6-8), and Cre^ER^*Rraga*^fl/fl^*Rragb*^fl/fl^*Tfeb*^fl/fl^ (n = 4-6) mice. Data in graphs represent mean ± SEM. ns, not significant. *p < 0.05, **p < 0.01, and ***p < 0.001, one-way ANOVA (C, E and K), two-way ANOVA (H, I and J).

**Figure S6.**
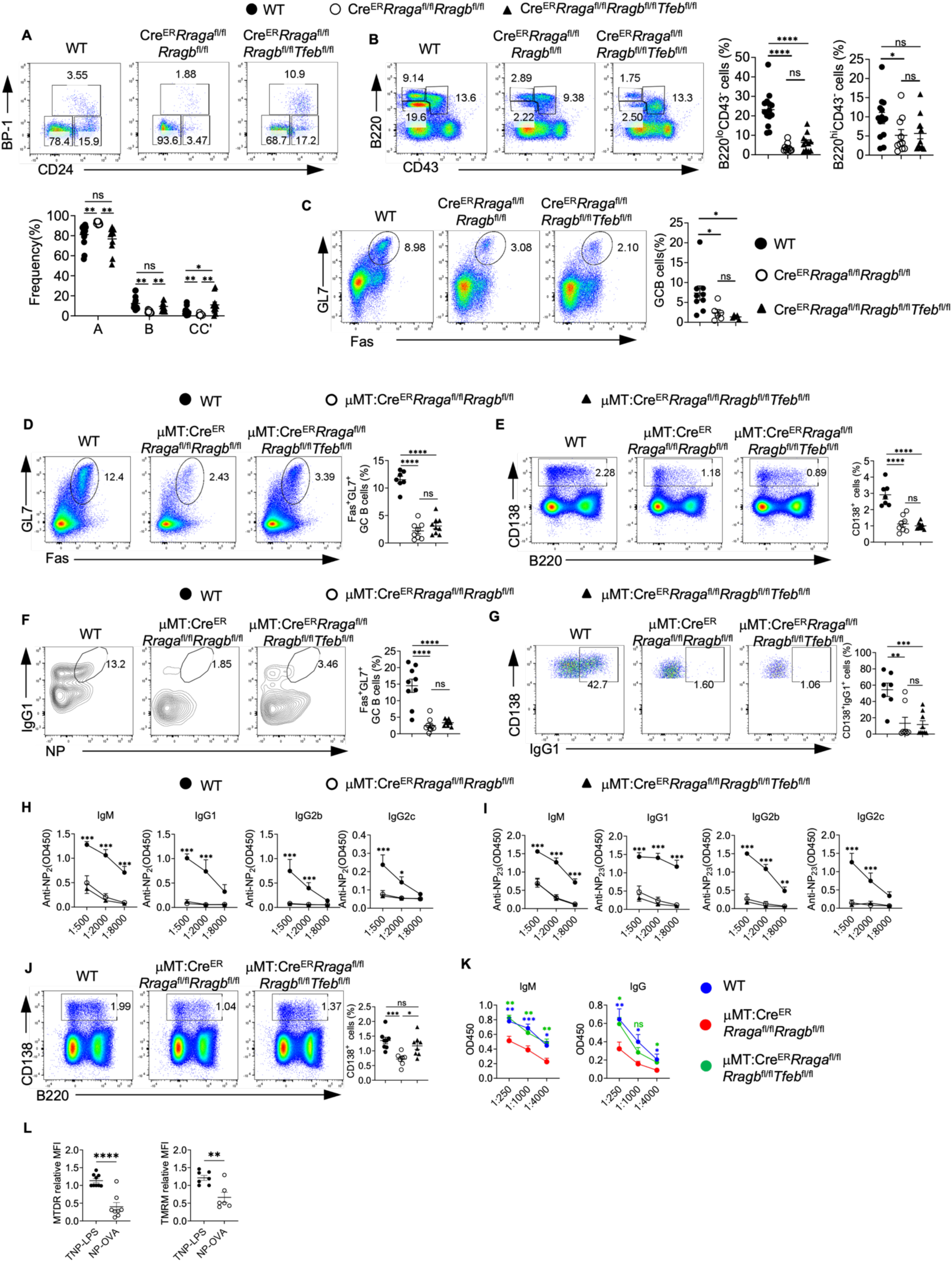
Abrogating TFEB rescues the deficiency in Rag-GTPases KO mice in context-dependent manner. (A-C)Tamoxifen was administered to animals intraperitoneally daily for 4 consecutive days. Mice were sacrificed and analyzed 7 days after the last injection. (A) Representative flow plots of BP-1 and CD24 expression in BM B220^+^CD43^+^IgM^−^ B cell precursors from WT (n = 14), Cre^ER^*Rraga*^fl/fl^*Rragb*^fl/fl^ (n = 10), and Cre^ER^Rraga^fl/fl^*Rragb*^fl/fl^*Tfeb*^fl/fl^ (n = 11) mice. Right, summary of the percentages of fraction A (CD24^−^BP-1^−^), fraction B (CD24^+^BP-1^−^), and fraction C/C′ (CD24^+^BP-1^+^) cells. (B) Representative flow plots of B220 and CD43 expression in BM lymphocytes from Cre^ER^*Rraga*^fl/fl^*Rragb*^fl/fl^ (n = 10), Cre^ER^Rraga^fl/fl^*Rragb*^fl/fl^*Tfeb*^fl/fl^ (n = 11), and WT (n = 14) mice. Right, summaries of the percentages of B220^lo^CD43^−^ and B220^hi^CD43^−^ cells. (C) Representative flow plots of GL-7 and Fas expression in lymphocytes from the Peyer’s patches of WT (n = 9), Cre^ER^*Rraga*^fl/fl^*Rragb*^fl/fl^ (n = 6), and Cre^ER^Rraga^fl/fl^*Rragb*^fl/fl^*Tfeb*^fl/fl^ (n = 5) mice. Right, summary of the percentages of GC (GL-7^+^Fas^+^) B cells. (D-I) Tamoxifen was administered to animals by oral gavage daily for 4 consecutive days. Mice were immunized intraperitoneally (100 mg NP-OVA/alum) 7 days after the last injection. (D) Representative flow plots of GL-7 and Fas expression on splenic B cells from the spleen of WT (n = 7), μMT:Cre^ER^*Rraga*^fl/fl^*Rragb*^fl/fl^ (n = 8), and μMT:Cre^ER^*Rraga*^fl/fl^*Rragb*^fl/fl^*Tfeb*^fl/fl^ (n = 9) immunized chimera mice. Right, summary of the percentages of GC (GL-7^+^Fas^+^) B cells. (E) Representative flow plots of CD138 and B220 expression in spleens from immunized WT (n = 7), μMT:Cre^ER^*Rraga*^fl/fl^*Rragb*^fl/fl^ (n = 8), and μMT:Cre^ER^*Rraga*^fl/fl^*Rragb*^fl/fl^*Tfeb*^fl/fl^ (n = 9) chimera mice. Right, summary of the percentages of CD138^+^ cells. (F) Representative flow plots of NP^+^IgG1^+^ GC B cells from the spleen of immunized WT (n = 7), μMT:Cre^ER^*Rraga*^fl/fl^*Rragb*^fl/fl^ (n = 9), and μMT:Cre^ER^*Rraga*^fl/fl^*Rragb*^fl/fl^*Tfeb*^fl/fl^ (n = 9) chimera mice. (G) Representative flow plots of splenic CD138^+^IgG1^+^ cells from immunized WT (n = 7), μMT:Cre^ER^*Rraga*^fl/fl^*Rragb*^fl/fl^ (n = 8), and μMT:Cre^ER^*Rraga*^fl/fl^*Rragb*^fl/fl^*Tfeb*^fl/fl^ (n = 9) chimera mice. (H) high-affinity NP-specific antibodies of all classes were measured using the sera of immunized WT (n = 6), μMT:Cre^ER^*Rraga*^fl/fl^*Rragb*^fl/fl^ (n = 8), and μMT:Cre^ER^*Rraga*^fl/fl^*Rragb*^fl/fl^*Tfeb*^fl/fl^ (n = 9) chimera mice. (I) Total NP-specific antibodies of all classes were measured using the sera of immunized WT (n = 6), μMT:Cre^ER^*Rraga*^fl/fl^*Rragb*^fl/fl^ (n = 8), and μMT:Cre^ER^*Rraga*^fl/fl^*Rragb*^fl/fl^*Tfeb*^fl/fl^ (n = 9) chimera mice. (J) Representative flow plots of splenic CD138^+^ cells from TNP-LPS immunized WT (n = 8), μMT:Cre^ER^*Rraga*^fl/fl^*Rragb*^fl/fl^ (n = 7), and μMT:Cre^ER^*Rraga*^fl/fl^*Rragb*^fl/fl^*Tfeb*^fl/fl^ (n = 8) chimera mice. (K) anti-TNP antibody titers in the sera of immunized WT (n = 8), μMT:Cre^ER^*Rraga*^fl/fl^*Rragb*^fl/fl^ (n = 7), and μMT:Cre^ER^*Rraga*^fl/fl^*Rragb*^fl/fl^*Tfeb*^fl/fl^ (n = 8) chimera mice. (L) WT mice were immunized with TNP-LPS or NP-OVA/alum, TMRM or MTDR on CD138^+^ cells were measured at day 9 (NP-OVA/alum) or 7 (TNP-LPS) post-immunization. Data in graphs represent mean ± SEM. ns, not significant. *p < 0.05, **p < 0.01, ***p < 0.001, and ****p < 0.0001, one-way ANOVA (B, C, D, E, F, G and J), two-way ANOVA (A, H, I and K), two-tailed Student’s t test (L).

